# The cysteine-rich exosporium morphogenetic protein, CdeC, exhibits self-assembly properties that lead to organized inclusion bodies in *Escherichia coli*

**DOI:** 10.1101/2020.07.09.196287

**Authors:** A. Romero-Rodríguez, S. Troncoso-Cotal, E. Guerrero-Araya, D. Paredes-Sabja

## Abstract

*Clostridioides difficile* is an obligate anaerobe spore-forming, Gram-positive, pathogenic bacterium, considered the leading cause of nosocomial diarrhea worldwide. Recent studies have attempted to understand the biology of the outer-most layer of *C. difficile* spores, the exosporium, which is believed to contribute to early interactions with the host. The fundamental role of the cysteine-rich proteins CdeC and CdeM has been described. However, the molecular details behind the mechanism of exosporium assembly are missing. The underlying mechanisms that govern exosporium assembly in *C. difficile* remain poorly studied, in part due to difficulties in obtaining pure soluble recombinant proteins of the *C. difficile* exosporium. In this work, we observed that CdeC was able to form organized inclusion bodies in the *E. coli* BL21 (DE3) pRIL strain filled with lamellae-like structures separated by an interspace of 5-15 nm; however, this lamellae-like organization is lost upon overexpression in *E. coli* SHuffle T7 strain with an oxidative environment. Additionally, DTT treatment of CdeC inclusion bodies released monomeric soluble forms of CdeC. Three truncated versions of the CdeC protein were constructed. While all the variants were able to aggregate forming oligomers that are resistant to denaturation conditions, TEM micrographs suggest that the self-organization properties of CdeC may be attributed to the C-terminal domain. Overall, these observations have important implications in further studies implicated in elucidating the role of CdeC in the exosporium assembly of *C. difficile* spores.

## Introduction

*Clostridioides difficile* is an obligate anaerobe spore-forming, Gram-positive, a pathogenic bacterium, considered the leading cause of nosocomial diarrhea worldwide (Burke & Lamont 2014, Sebaihia et al 2006). *C. difficile* infection (CDI) symptoms range from mild diarrhea to pseudomembranous colitis and toxic megacolon that can be life-threatening. The CDI mortality rate is 5% and may increase to 20% in more severe cases (Evans & Safdar 2015). *C. difficile* spores are allowed long-term persistence outside the host, facilitating transmission from host to host and provides recalcitrance against anti-infective stressors (Deakin et al 2012). Therefore, spores are the vehicle for dispersion, transmission, and persistence. Recent reports have estimated the overall incidence rate of CDIs for patients of all ages was 2.24 per 1000 admissions per year (Balsells et al 2019). The increased incidence and severity of CDI has led to a significant economic burden on healthcare systems due to the costs associated with treatment, and extended stays of patients in hospital (Collins & Auchtung 2017, Elliott et al 2017).

Important in understanding how this pathogen interacts with the host and for the development of counter measurements, is the spore surface. Recent studies have attempted to understand the biology of the outer-most layer of *C. difficile* spores, the exosporium, which is believed to contribute to early interactions with the host (Calderon-Romero et al 2018, Mora-Uribe et al 2016, Phetcharaburanin et al 2014). In a proteomic gel-free study, Díaz-González et al. (2015) demonstrated that the outer-most exosporium layer of *C. difficile* spores has a total of 184 proteins, importantly 7 of which have been previously characterized as spore coat and/or exosporium proteins, and 13 have been identified as uncharacterized proteins unique to *C. difficile* (Diaz-Gonzalez et al 2015). The exosporium in *B. cereus* group has a basal layer, which contains cysteine-rich proteins, such as CotY and ExsY, that assemble into a porous hexagonal lattice and a hair-like protrudes from the basal layer (Jiang et al 2015, Stewart 2015, Terry et al 2017). In the *B. cereus* group, the hair-like nap is formed mainly by radial projections of a high collagen-like glycosylated protein, BclA, which is anchored in the basal layer (Stewart 2015). A group of structural proteins, which participate in the assembly of the exosporium of the *B. cereus* group, such as CotE, CotY, ExsA, ExsB, ExsY and ExsM (Bailey-Smith et al 2005, Boydston et al 2006, Giorno et al 2007, McPherson et al 2010) has been described. In *C. difficile*, these proteins are absent in the exosporium proteome, but orthologous of the BclA family have been identified. Besides, the cysteine-rich proteins CdeC and CdeM have been identified (Alves Feliciano et al 2019, Barra-Carrasco et al 2013, Calderon-Romero et al 2018, Diaz-Gonzalez et al 2015), which have a predominant role in the exosporium function and assembly(Antunes et al 2018, Calderon-Romero et al 2018).

The underlying mechanisms that govern exosporium assembly in *C. difficile* remain understudied, in part due to difficulties in obtaining pure soluble recombinant proteins of the *C. difficile* exosporium. Recently, we evaluated the effect of different heterologous *Escherichia coli* strains to increase soluble levels of exosporium proteins of *C. difficile* spores (Brito-Silva et al 2018). Their results indicate that optimum soluble expression conditions may vary between 21, 30, and 37°C, depending on the protein, and at least CdeC, BclA1, and BclA3, required strains that provided an oxidative environment such as *E. coli* SHuffle T7 for increased soluble levels. However, whether these observations can be applied to additional cysteine-rich exosporium proteins remains unclear.

In this work, while attempting to expand these findings to additional cysteine-rich proteins, CdeA and CdeM, we observed that only CdeC was able to form organized inclusion bodies in the *E. coli* BL21 (DE3) pRIL strain, which was filled with lamellae-like structures with an interspace of 5-15 nm. Moreover, we observed that these self-assembly properties were independent of the expression vector being used, temperature, and time of expression. However, upon overexpression in *E. coli* SHuffle T7 strain, the increased oxidative environment hampered CdeC’s ability to self-organize. Moreover, DTT treatment allows the release of soluble CdeC. To expand our analysis, three truncated versions of CdeC were constructed, and the SDS-PAGE analysis revealed that all CdeC variants could aggregate forming oligomers that are resistant to denaturation conditions and TEM micrographs analysis indicate that the self-organization properties of CdeC may be attributed to the C-terminal domain of the protein. These observations have important implications in further studies related to the role of CdeC in exosporium-assembly of *C. difficile* spores.

## Material and methods

### Bacterial strains, plasmids, and media

*Escherichia coli* BL21 (DE3) pRIL and SHuffle T7 (New England Biolabs, USA) strains were routinely grown in Luria Bertani (LB). The expression of recombinant proteins in *E. coli* was performed in LB broth supplemented with 0.5% of glucose (LBG) (21 or 37°C at 2 x *g*). The detailed description of the genetic background and relevant characteristics of each strain is provided in Table S1.

### Construction of overexpression plasmids

Phenol-chloroform extracted genomic DNA from *C difficile* R20291 (Accession No. FN545816.1) strain was used as a template for the amplification of *cdeC* (CD0926), *cdeM* (CD1478) and *cdeA* (CD2262) genes and the truncated variants of *cdeC*. All amplifications were carried-out with Phusion Hot Start High-Fidelity DNA polymerase (Thermo Scientific, USA). The resulting amplicons were cloned in plasmid pET22b and (or) pETM11. CdeC was cloned by the traditional restriction-ligation method in the pETM11 vector between N*co*I and X*ho*I restriction sites, while the construction in pET22b was done between the N*de*I and X*ho*I sites. The truncated version of CdeC was cloned in pET22b, as described before. CdeM and CdeA were cloned in pETM11 by Gibson assembly between N*co*I and X*ho*I sites. Sites (Gibson Assembly^®^ Cloning Kit, New England Biolabs). All primers are listed in Table S2. The maps of the resulting plasmids are shown in Table S1. All the constructs were verified by DNA digestion and Sanger sequencing.

### Overexpression of CdeA, CdeC, CdeM, and truncated variants of CdeC-6His fusions

Plasmids pDP339, pARR10 or pARR19, pARR21 and pARR22, pARR20, pARR7 and pARST1 containing *cdeC (C. difficile* strain 630, CD1067), *cdeC (C. difficile* strain R20291, CD0926)*, cdeM, cdeA,* and the *cdeC* variants *cdeC_1-100_, cdeC_1-214_* and *cdeC_206-405_*, respectively, were transformed into *Escherichia coli* strains BL21(DE3) pRIL. Also, plasmid pARR19 was transformed into *E. coli* strains BL21(DE3) pRIL and SHuffle T7 (Müllerová et al 2009). Transformed cells were used to inoculate 5 ml LBG medium (containing 50 μg ml^-1^ kanamycin or 150 μg ml^-1^) and were grown overnight at 37°C and 2 x *g.* The overnight cultures (5 ml) were used to inoculate 600 ml of production medium. Cultures were grown to optical densities between 0.7 and 0.9 at 600 nm, after which the temperature maintained at 37°C or reduced to 21°C. The cultures were induced with 0.5 mM of isopropyl-β-D-thiogalactoside (IPTG). Incubation was continued for 16 h, and at 37°C and 2 x *g,* the cells were harvested by centrifugation at 5853 x *g* for 15 min. The cell pellets were stored at −80°C.

Next, cells were harvested and first resuspended in 500 μL of soluble lysis buffer, 20mM Tris-HCl (pH 7.8), 500mM NaCl, 5mM imidazole, 0.5–0.8% Triton X-100 and 0.1 mM Protease inhibitors (Thermofisher, U.S.A.). Resuspended cultures were sonicated at the output of 12 W for six bursts of 15 s, separated by 15 s of cooling on ice-cold water. Lysates were centrifuged at 18,625 g for 10 min, and the supernatant was saved as the soluble fraction, and the pellet was subsequently resuspended in 500 μL of insoluble lysis buffer, 20mM Tris-HCl (pH 7.8), 500mM NaCl, 5mM imidazole and 8 M urea. Resuspended lysates were sonicated at the output of 21 W for six bursts of 30 s, separated by 30 s of cooling on ice-cold water and saved as the insoluble fraction. Soluble and insoluble fractions were further analyzed by SDS-PAGE and Western blot, as described below.

### SDS-PAGE and western blot analysis of recombinant protein CdeC, CdeM, CdeA and truncated variants of CdeC

Fraction soluble and insoluble of recombinant CdeC, CdeM and CdeA were mixed with 2× SDS-PAGE loading buffer and then were boiled for 5 min, and we loaded 2 μg of total protein per lane. Samples were electrophoresed in SDS-PAGE gels (15% acrylamide). Some gels were directly stained with Coomassie brilliant blue, and in others, proteins were transferred to nitrocellulose membrane (Bio-Rad, U.S.A.), blocked 1 hour at room temperature with 5% skim milk in Tween-Tris-buffered (TTBS) (pH 7.4). The blocked membranes were probed overnight at 4°C temperature with a 1:5,000 dilution of mouse anti-6His antibodies (Rockland Immunochemicals Inc., USA or Thermo), rinsed three times with TBS-0.1% (v/v) Tween 20 and incubated for 1h at room temperature with 1:10,000 dilution of anti-mouse IgG-horseradish peroxidase conjugate (HRP) (Rockland Immunochemicals Inc., USA). Both primary and secondary antibodies were incubated in the presence of 5% milk (Sigma-Aldrich). HRP activity was detected with a chemiluminescence detection system (Li-COR Imaging System) by using Clarity chemiluminescent detection system HRP substrate (Bio-Rad). Each Western blot also included 3 μL of a Page Ruler Plus pre-stained protein ladder (Thermo, USA).

### Purification of inclusion bodies

Recombinant CdeC was expressed in *E. coli* BL21 (DE3) pRIL according to the following conditions: the expression was induced with 0.5 mM IPTG for 16 hours at 37°C and with 2 x *g* agitation in LBG medium. Cells were concentrated by centrifugation at 5853 x *g* for 15 min at 4°C, and the inclusion bodies were isolated as described below. The pellet was resuspended with 30-35 mL of 10 mM EDTA in PBST 1X buffer per liter of overexpression culture. The cells were then lysed with 10 mg/mL lysozyme for 1 hour at 37°C. After that, cells were sonicated at 12 Watts for 15 s on ice and centrifuged at 5853 x *g* for 45 min at 4°C. The supernatant obtained was discarded, and the pellet was washed two times with 2% Triton X-100 in PBS 1X buffer by centrifugation at 5853 x g for 20 min at 4°C. The lysate cells were sonicated at a power of 12 Watts for 15 s on ice and passed through a 45% Nicodenz gradient for 50 min at 5853 x *g* at 4°C to separate the cells and inclusion bodies. Finally, the inclusion bodies were washed twice with 0.1 mM PMSF in 1X PBS buffer by centrifugation at 5853 x *g* for 20 min, adjusted to optical densities of 0.2 at 600 nm in a volume of 20 μL and stored at −80°C.

### Bioinformatic analysis

1835 publicly available *C. difficile* genomes were downloaded from GenBank, and their MLST and clade were determined in the PubMLST database (Table S3). The amino acid sequence of CdeC was searched using tBLASTn v 2.9.0+. The nucleotide sequences of the CDS corresponding to the *cdeC* gene were extracted using as threshold hits with at least 90% coverage concerning the reference gene of the *C. difficile* 630 strain leaving all parameters by default. The genes found are in the supplementary Table S4 In order to make a rooted phylogenetic tree explaining the evolutionary history of CdeC, the NCBI RefSeq database was searched using the same strategy, leaving out all hits with organisms that were described as *C. difficile* (tax id: 1496) and that had a coverage of less than 90% to the reference gene of the *C. difficile* 630 strain. The genes found and the species to which it belongs are in the supplementary Tables S5 and S6 and Table S4. The files were concatenated and clustered into single alleles using CD-HIT-EST v4.8.1. The single alleles are in Table S7. These single alleles were aligned using the AlignTranslation function of the R DECIPHER package v2.16.1 with the default parameters. The multiple sequence alignment is found in the supplementary Table S8 (nucleotide) and Table S9 (amino acid). To perform the phylogenetic inference, the GTR+I+G substitution model was determined by the AIC and BIC methods using the jModelTest v2.1.10 program on nucleotide alignment Table S8 Phylogenetic inference was performed using the Bayesian method with the program BEAST v1.10.4, using for this purpose 6 independent chains of 10 000 000 states sampling every 1000 states. Valid sample size values were >200 for all parameters, and convergence and mixing were assessed using Tracer v1.7.1. Log and tree files were summarized with tree annotator v2.4.8 using 10 % burning, along with maximum clade credibility and node heights at the median. To complement the phylogenetic inference, a second inference was made using the Maximum Likelihood method with the RAxML v8.2.12 program, indicating that the outgroup taxa of the analysis are from the sequences that do not belong to the *C. difficile* species. Topologies comparison of the two trees was examined using the web service phylo.io, and the visualization of the phylogenetic trees was done in itol.embl.de. Also, because cysteines are amino acids that can command the transition from disorder to order as a result of changes of redox potential in the environment redox-sensitive disordered regions were evaluated on all single allele proteins using (Mészáros et al 2018) webserver. To predict the contact map of CdeC, we have used the program DeepMetaPSICOV 1.0 (Kandathil et al 2019) server under default settings

### DTT treatment of CdeC inclusion bodies

The isolated CdeC inclusion bodies were treated with different concentrations of dithiothreitol (DTT; Thermo Scientific, USA), a disulfide reducing agent, during different incubation times. CdeC inclusion bodies partially purified and adjusted to optical densities of 0.2 at 600 nm in a volume of 20 μL were treated with 500 mM, 1, and 2 M of DTT during 30, 60, and 120 min of incubation at 37°C. The effect of treatment of CdeC inclusion bodies with 2M concentration of DTT during different incubation times was evaluated by SDS-PAGE, western blot, and immunofluorescence.

### Immunofluorescence of overexpression recombinant cysteine-rich proteins and inclusion bodies of CdeC with or without DTT-treatment

The immunofluorescence analysis was performed to the overexpression of recombinant CdeC, CdeM, and CdeA proteins in *E. coli* and the partially purified CdeC inclusion bodies treated and not treated with DTT. For this purpose, recombinant CdeC, CdeM, and CdeA proteins were overexpressed in *E. coli* strain BL21 (DE3) pRIL, as described in the corresponding section An aliquot of 1 mL, was taken from both IPTG-induced and non-IPTG-induced cultures and concentrated by centrifugation at 4107 x *g* for 8 min at 4°C. Before the immunofluorescence analysis, *E. coli* cells were fixed in the LBG medium supplemented with 3% paraformaldehyde (PFA) and incubated for 10 min at room temperature and 30 min on ice. The *E. coli* cells were washed 3 times with PBS 1X by centrifugation 4107 x *g* for 8 min at 4°C and resuspended in GTE buffer (glucose 50 mM, EDTA 10 mM, and Tris 20 mM pH 7.5) supplemented with 0.7 mg/mL lysozyme. In the case of the partially purified CdeC inclusion bodies not treated with DTT, the immunofluorescence analysis was performed directly. In contrast, the inclusion bodies of CdeC treated with DTT were treated as described in the DTT Treatment section and washed 3 times with PBS 1X by centrifugation 5853 x *g* for 20 min at 4°C to remove excess DTT.

For immunofluorescence analysis, droplets of 2 μL samples were added to the poly-L-lysine pre-treated coverslips and dried for 10 min at 37°C. The samples were fixed with 4% PFA during 15 min at room temperature, except for *E. coli* cells which were previously fixed, and were washed three times with PBS 1X and once with H_2_O Milli Q. It was blocked with 1% BSA on PBS 1X for 1 h at room temperature in a humid chamber. Then, it was incubated with primary antibody α-Mouse 6xHis (Thermo Scientific, USA) in a 1:500 dilution in PBS 1X with BSA 1% for 1 h at room temperature in a humid chamber and 3 washes were performed with PBS 1X and one with H_2_O Milli Q. It was incubated with secondary antibody α-Mouse Alexa Fluor 488 (Abcam, UK) in a 1:400 dilution in PBS 1X with BSA 1% during 1 h at room temperature in a humid chamber and darkness. Later, three washes were made with PBS 1X and one with H_2_O Milli Q. Finally, the coverslips were dried for 10 min at 37°C, mounted with 3 μL of Dako, dried for 30 min at 37°C and stored at 4°C until observed under Olympus BX53 fluorescence microscope.

### Transmission electronic microscopy

Recombinant proteins CdeC, CdeM, CdeA, and truncated variants of CdeC were expressed in *E. coli,* as described in the corresponding section. The *E. coli* cells were concentrated by centrifugation at 5853 x *g* for 15 min and fixed with 3% glutaraldehyde on a 0.1M cacodylate buffer (pH 7.2), incubated overnight at 4°C, and stained for 30 min with 1% tannic acid. Samples were further embedded in a Spurr resin (Paredes-Sabja et al 2012). Thin sections of 90 nm were obtained with a microtome, placed on glow discharge carbon-coated grids, and double lead stained with 2% uranyl acetate and lead citrate. Spores were analyzed with a Philips Tecnai 12 Bio Twin microscope at Unidad de Microscopía Avanzada in Pontificia Universidad Católica de Chile.

For the measurements of the distance between each lamination, at least five independent inclusion bodies were divided in their axial and longitudinal axis, giving four sections of each inclusion body. From each section, five lamellae-like were measured, giving a total of 20 measurements per inclusion bodies. The results were plotted in a frequency distribution graph. The data were fit to a Gaussian curve using the Graph Pad Prism 8 software.

## Results

### Cysteine-rich protein multimerizes during heterologous expression

In a recent study, it was shown that the cysteine-rich protein CdeM exists not just as a monomer (predicted the molecular weight of 19 kDa) but also as several forms, including species of 25 and 60 kDa presumably stabilized by the formation of disulfide bonds. CdeC was localized to the spore as a high-molecular-mass complex (Barra-Carrasco et al 2013). Furthermore, upon heterologous expression, in *E. coli, CdeC* is readily detectable as a 42- and 84-kDa monomeric and dimeric immunoreactive species (Barra-Carrasco et al 2013). Brito *et al.* (Brito-Silva et al 2018) described optimized conditions for overexpression of exosporium proteins CdeC, BclA1, BclA2, and BclA3 in *E. coli* strain as a heterologous host. However, this work did not assess the formation of high molecular stable-complexes or aggregates.

Here, we expanded our previous work to look for the formation of complexes upon heterologous overexpression of the cysteine-rich proteins CdeA, CdeC, and CdeM, all of which were found in the exosporium proteome of *C. difficile* spores (Diaz-Gonzalez et al 2015). To gain information on the multimerization in soluble and insoluble fractions, the CdeA, CdeC, and CdeM genes coding were cloned in the pETM11 vector and expressed in *E. coli* BL21(DE3) pRIL as described. Under the tested conditions (growth on LBG at 37°C and induction with 0.5 mM IPTG), SDS-PAGE analysis of soluble and insoluble fractions did not demonstrate clear abundant bands corresponding to the molecular weights of the CdeA (11,4 kDa), CdeC (44,7 kDa), and CdeM (19,2 kDa) (Fig. 1). For CdeC, Western blot analysis of the insoluble fraction revealed at least three anti-His immunoreactive bands, probably multimerized forms (Fig. 1) of apparent molecular weights of 40, 60, and 130 kDa. On the other hand, Western blot analysis of the soluble fraction revealed an immunoreactive of an apparent molecular weight of 60 kDa, presumably, the monomeric form of CdeC. Also, Western blot analysis of the soluble and insoluble fraction showed a molecular weight band of approximately 35 kDa, which likely c a degradation product of CdeC.

**Fig. 1.**
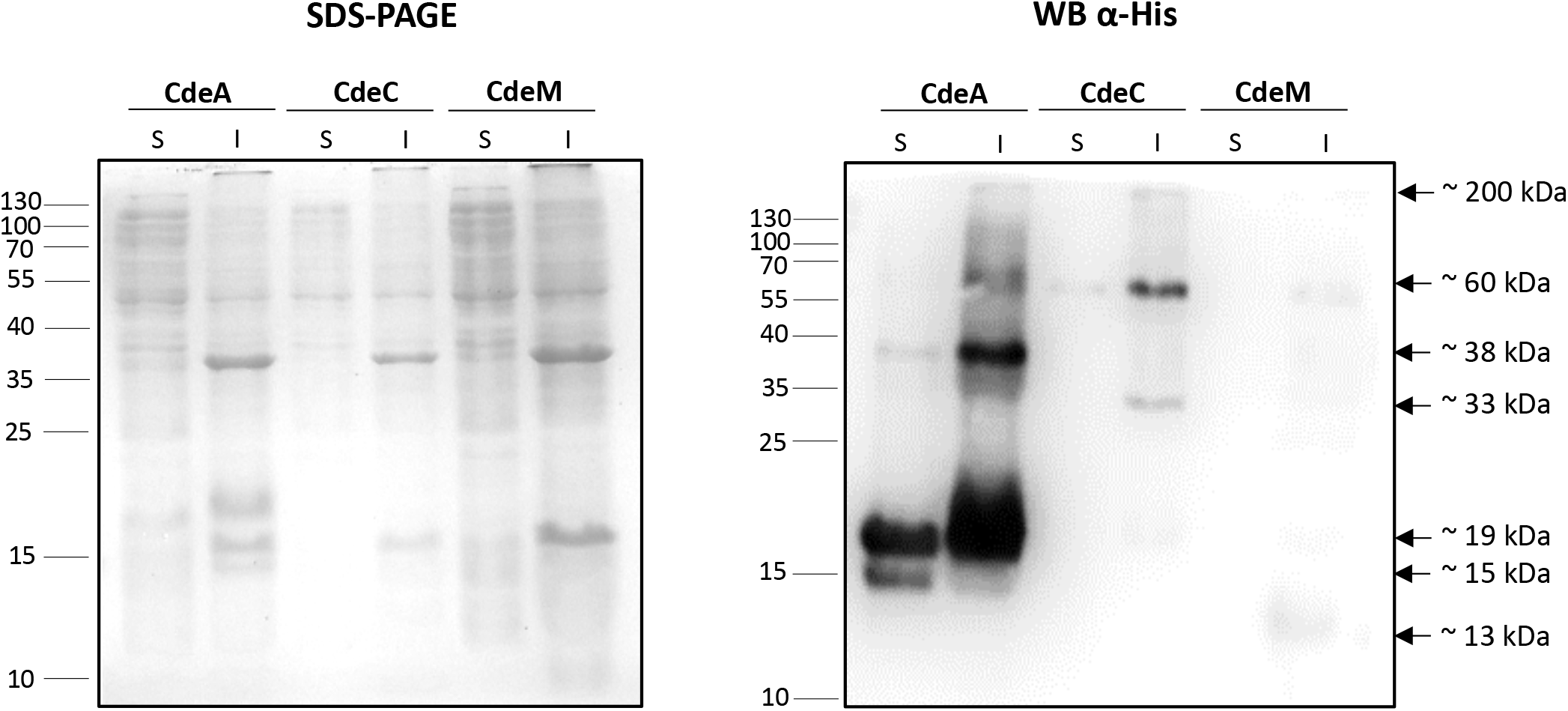
Heterologous overexpression of the cysteine-rich proteins CdeA, CdeC, and CdeM in the soluble (S) and insoluble (I) fractions of *E. coli* lysates. Recombinant proteins were expressed in *E. coli* BL21 (DE3) pRIL carrying plasmids pARR22, pARR10, and pARR21, respectively, and induced with 0.5 mM IPTG for 16 h at 37°C. The cells were disrupted in soluble or insoluble lysis buffer, electrophoresed in 15% SDS-PAGE, and His-tagged immunoreactive proteins detected by Western blot analysis as described in the Methods section. Each lane was loaded with 2 μg of protein lysate. Molecular mass (kDa) markers are indicated at the *left side* of the panels, and molecular mass of the detected His-tagged immunoreactive bands are indicated at the *right side* of the panels.

For the CdeM, the Western blot analysis of the insoluble fraction showed at least three immunoreactive bands of approximately 19, 60, and 130 kDa likely multimerized forms of CdeM (Fig. 1). The Western blot analysis of the soluble fraction revealed an immunoreactive band of an apparent molecular weight of 19 kDa, the putative monomeric form of CdeM.

In the case of CdeA, Western blot analysis of the insoluble fractions revealed at least three immunoreactive bands (Fig. 1) with apparent molecular weights 19, 38, and 60 kDa. The immunoreactive bands probably are multimerized forms of CdeA. For the Western blot analysis of the soluble fraction revealed two immunoreactive bands of 19 and 38 kDa. Presumably, the multimeric form with a molecular weight of approximately 19 kDa could be to the monomeric form of CdeA. These results suggest that the three cysteine-rich proteins from the exosporium of *C. difficile* form multimers upon heterologous expression in *E. coli*. The multimers are found in the insoluble fraction, while the monomeric forms are found in the soluble fraction. Interestingly, the multimers formed by these proteins are highly stable since they are still observed under the harsh conditions in the buffer for the extraction of the insoluble fraction.

### Cysteine-rich proteins CdeC and CdeM, but not CdeA, are found in large inclusion bodies

Due to the weak soluble expression of CdeC and CdeM proteins, cells overexpressing these proteins were observed under the microscope, and the formation of large inclusion bodies was detected for the cells expressing CdeC and CdeM (Fig. 2A). These aggregates or inclusion bodies (IBs) can be observed as large refractive bodies that are predominantly located at one cell pole. To confirm that those large structures correspond to the heterologous proteins, immunofluorescence against the His-tag was done. The signal localizes in the round structures observed in the contrast phase (Fig. 2B). To gain further information about the putative structural features of the inclusion bodies, cells expressing the recombinant proteins were observed by transmission electron microscopy (TEM). As expected, in the cytoplasm of the cells expressing CdeA, no clear inclusion bodies were detected (Fig. 2C). Instead, in the case of CdeC and CdeM expressing cells, large rounded inclusion bodies were observed. Surprisingly, TEM micrographs revealed that in the case of CdeC, the ultrastructure of the inclusion bodies appeared as organized lamellae-like structures (Fig. 2C). These results suggest that CdeC can form organized ultrastructure, not just aggregates as CdeM.

**Fig. 2.**
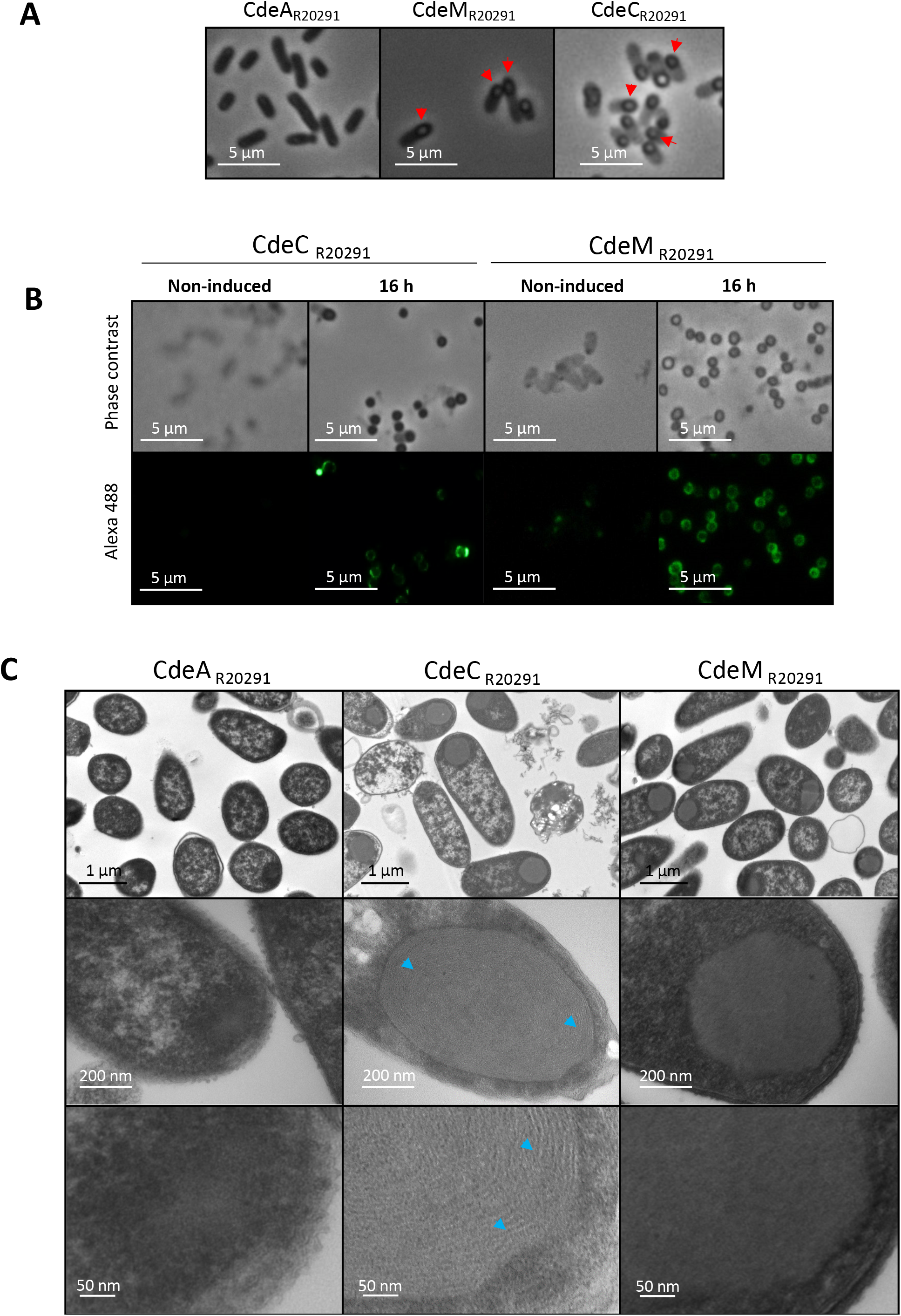
Microscopy analysis of *E. coli* cells expressing the cysteine-rich proteins CdeA, CdeC, and CdeM. Recombinant proteins were expressed in *E. coli* BL21 (DE3) pRIL carrying plasmids pARR22, pARR10, and pARR21, respectively, and induced with 0.5 mM IPTG for 16 h at 37°C in LBG. A) Samples were withdrawn from the cultures at 16 h and observed under contrast phase microscopy. CdeC and CdeM examples of inclusion bodies are indicated with a red arrowhead. B) Immunofluorescence analysis of *E. coli* BL21 (DE3) pRIL cells carrying plasmids pARR22, pARR10, and pARR21 without inductor or after induction for 16 h. α-6xHis was used as the first antibody while the secondary antibody was conjugated with Alexa-488 fluorophore. C) Electron microscopy analysis of *E. coli* BL21 (DE3) pRIL cells expressing CdeA, CdeC, or CdeM (plasmids pARR22, pARR10, and pARR21, respectively). The samples were taken at 16 h post-induction, as previously mentioned. Representative micrographs of several *E. coli* cells are shown in the upper panel. The middle panel shows selected individual cells. Lower panel is a magnification of individual cells. When corresponding, lamellae-like formation in CdeC is indicated with blue arrowheads.

### Inclusion bodies of CdeC, but not CdeM, exhibit conserved organized structure

As mention before, the overexpression of CdeCR_20291_ in the vector pETM11 resulted in the generation of inclusion bodies. Nevertheless, unlike the inclusion bodies of CdeM, those of CdeC showed an organized lamellae-like ultrastructure. Besides overexpression in pETM11, CdeC was also cloned and expressed in the vector pET22b with no differences in the lamellae ultrastructure (Fig S1A). Also, the overexpression was tested at 21°C, and 37°C, no differences in the lamellae-like structure were evidenced (Fig. S1B), indicating that this is temperature independent.

The CdeC cysteine-rich proteins are highly conserved in Peptostreptococcaceae family members, and, at least in the epidemically relevant R20291 strain, it is essential for morphogenesis of the exosporium layer and spore resistance (Calderon-Romero et al 2018). We expand the phylogenetic analysis previously reported (Calderon-Romero et al 2018); for this, the analysis of *C. difficile* genomes began with 1835 MLST assigned genomes (Table S1). From these, 1833 alleles of *cdeC* were extracted (Table S4). Besides, 16 alleles of *cdeC*-like proteins with at least 90% coverage from the family Peptostreptococcaceae (Clostridium cluster XI) were added to extend the analysis to other species and to discriminate amino acids at positions preserved throughout evolution. The assemblies with the access codes were described in Table S2. To reduce the dataset without losing the diversity of the *cdeC*-like genes, they were clustered in 100% identity alleles. The information on the clusters obtained and the single alleles are described in tables S3 and S4. The multiple alignments (Fig. S2) of the CdeC shows interesting shared motifs along all the genes evaluated, which could be critical in the functionality of the protein. It should be noted that CdeC belonging to *C. difficile* are very homogeneous among themselves where the minimum identity at the amino acid level is greater than 90%, so an analysis without outgroups would not account for indispensable structural domains of CdeC. Phylogenetic reconstruction using two different methodologies were topologically identical, which suggests a robust phylogenetic analysis (Fig. S3).

According to Calderon-Romero et al. (2018), the sequence of CdeC contains several sequence motifs (Fig. 3A): i) in the N-terminal domain (NTD) two motifs of unknown function were identified (i.e., KKNKRR and three consecutive histidine residues); ii) a sequence of 3 consecutive histidine residues near the NTD; iii) in the central region, a 6 NPC repeat followed by two CCRQGKGK repeats; and iv) cysteine-rich sequence CNECC at the C-terminal domain (CTD). This analysis was performed with the amino acid sequence of CdeC from strain 630. However, upon comparing the sequences of CdeC between strains R20291 and 630, we observed at least six amino acid substitutions (Fig. 3A). However, those changes are likely conservative substitutions, as in the case of aspartate 57 (negatively charged; polar; hydrophilic) found in CdeC_630_ while in the same position glutamate (negatively charged; polar; hydrophilic) residue is present in CdeCR_20291_. Probably the most significant change, due to the chemical properties of the amino acids, is the presence of serine 229 (polar, non-charge) in CdeC_630_ while in the same position in CdeCR_20291_, the amino acid sequence includes an alanine (non-polar, aliphatic).

**Fig. 3.**
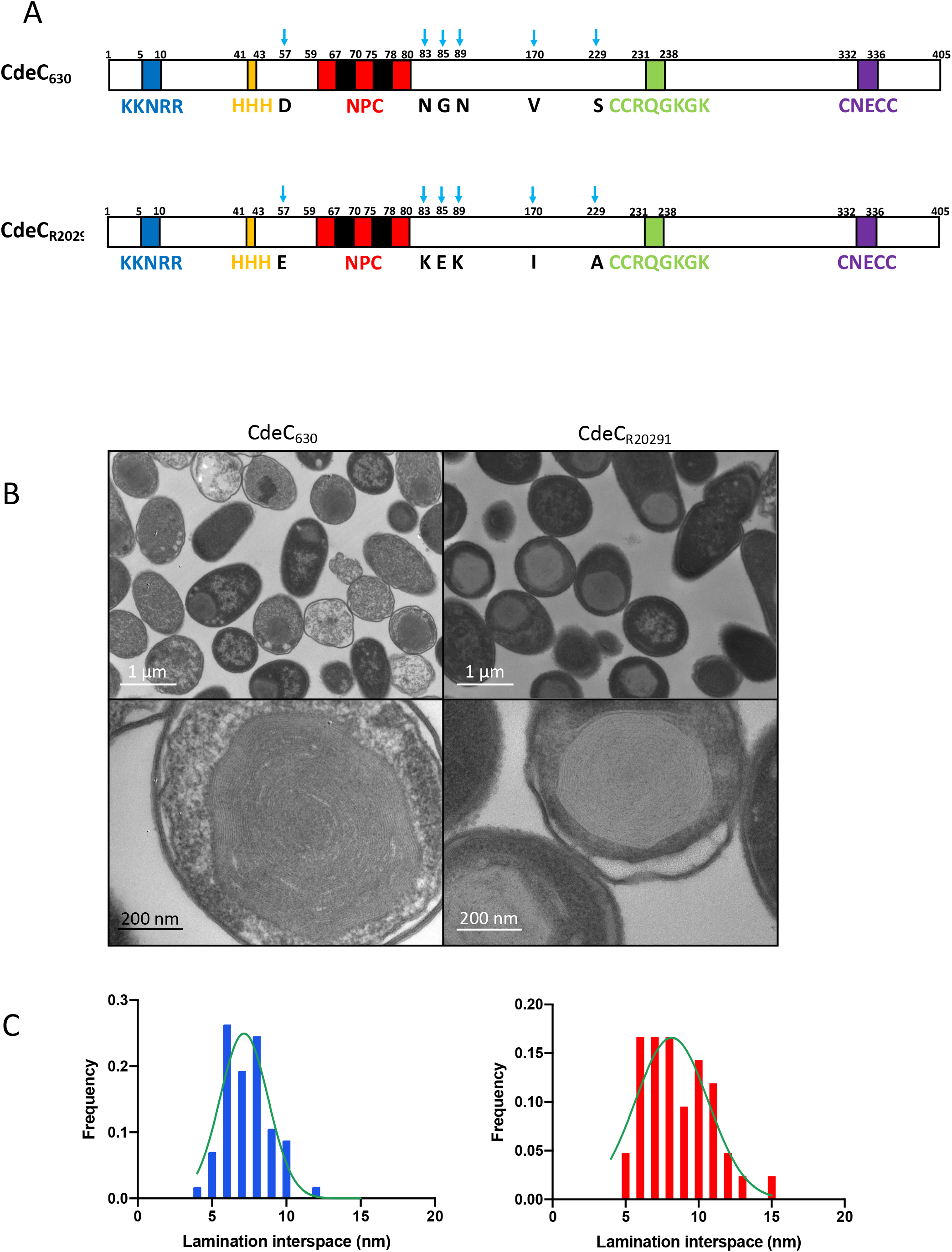
Amino acid conservation and ultrastructural organization of CdeC. A) Schematic representation of the CdeC sequence of *C. difficile* strains 630 and R20291, each amino acid substitution is indicated by a blue arrow B) *E. coli* BL21 (DE3) pRIL carrying plasmids expressing CdeC from *C. difficile* 630 (pDP339) or CdeC from R20291(pARR19), were induced with 0.5 mM IPTG for 16 h at 37°C. After induction time, samples were retrieved from cultures and subjected to TEM analysis. Representative micrographs of several thin sections of *E. coli BL21* (DE3) pRIL expressing CdeC from *C. difficile* 630 or *C. difficile* R20291 are shown in the upper panel. The lower panel shows selected individual cells. C) The distance between lamellae-like of at least ten sections of the inclusion bodies showing clear lamellae-like structure was measured, and the frequency distribution was plotted as a function of lamination interspace. The frequency lamination interspace of CdeC strain 630 (blue) and CdeC strain R20291 (red) is shown.

Consequently, we asked whether the formation of the lamella is a feature that is conserved in CdeC. To address this, we compared the ultrastructure of CdeC_630_ and CdeC_R20291_ from strains 630 and R20291, which belong to clades ST54 MLST clade1 and ST1 MLST clade 2, respectively. When observed in TEM, cells overexpressing CdeC from R20291 or 630, the formation of lamellae-like was present in both strains (Fig. 3B). Consequently, the differences in those amino acid residues seem to be irrelevant for the self-organization of CdeC when expressed in the heterologous host, *E. coli.* Furthermore, we measure the distance between each lamination in individual inclusion bodies. For CdeC_630_, the distance between each lamellae-like structure varied in the range of 5-12 nm, while for CdeC_R20291_, the distances were in the range of 5-15 nm. However, distances between lamellas with higher frequencies were 7 and 8 nm (Fig. 3C). These observations demonstrate that the properties of CdeC to form these self-assembled structures with well-defined laminations spans across *C. difficile* clades. However, the molecular basis of driving self-organization that leads to supramolecular structures is a matter of further studies.

### Effect of highly oxidizing cytoplasmic environment on the ultrastructure of CdeC inclusion bodies

Several covalent cross-links, such as ε-(γ-glutamyl)-lysyl isopeptide bonds are involved in stabilizing *the Bacillus subtilis* spore architecture (Fernandes et al 2019, Kobayashi et al 1998). Considering the abundant cysteine-residues in the CdeC amino acid sequence, it is feasible to the hypothesis that, at least part of the CdeC-self-assembly lamellae-like, may be impacted by the formation of disulfide bonds, as is the case in some of the spore coat cysteine-rich protein from *B. subtilis* (Jiang et al 2015). One approach to indirectly test the effect of the redox environment on the formation of the lamella is to use *E. coli* SHuffle strain. This strain has an oxidizing cytoplasm, which contributes to the formation of stable disulfide bonds between cysteine-residues of cytoplasmic proteins (Lobstein et al 2012). Further, this strain was also engineered to express DsbC in the cytoplasm, which isomerizes (rearranges), disulfide bonds to their native states. (Lobstein et al 2012). During the oxidative folding of proteins, disulfide bonds are likely to form between those cysteine residues that are most proximal in the amino acid sequence. However, for the proper fold of the protein, those nascent disulfide bonds must rearrange (isomerize) to the half-cystine (the group formally derived by the removal of a hydrogen atom from the thiol group on the side chain of a cysteine) pairings of the native conformation (Kersteen et al 2005).

Consequently, to test the influence of an oxidizing cytoplasmic environment in the formation of multimeric species of CdeC, lysates prepared from *E. coli* Bl21 (DE3) pRIL and *E. coli* SHuffle T7 strains expressing CdeC were tested. Western blot analysis showed the formation of different multimeric forms of CdeC in strains, *E. coli* Bl21 (DE3) pRIL and from *E. coli* SHuffle T7 (Fig. 4A). First, in *E. coli* BL21 (DE3) pRIL, we observed that soluble CdeC was present as 38 and 60-kDa immunoreactive species, while insoluble fractions revealed the presence of four immunoreactive species of 33, 38, 60 and 130 kDa species. However, upon CdeC overexpression in *E. coli* SHuffle T7, no immunoreactive CdeC was detectable in soluble *E. coli* lysates. By contrast, two immunoreactive species of 60 and 130-kDa were evidenced in insoluble *E. coli* SHuffle T7 lysates. These results suggest that the multimeric forms with a molecular weight of approximately 33 and 38 kDa could be a degradation product of the 44-kDa CdeC, processing that does not occur in a more oxidative cytoplasm such as that of *E. coli* SHuffle T7. Though, CdeC oligomerization was not affected by the oxidative state of the cytoplasm. Next, we asked whether an oxidative cytoplasm affects the formation of inclusion bodies with lamellae-like upon analyzing through transmission electron micrographs. As aforementioned, the inclusion bodies from *E. coli* BL21 (DE3) pRIL displayed an organized structure formed by lamellae-like, while the E. *coli* SHuffle inclusion bodies lost the lamellae-like structure and were observed as amorphous aggregates. Overall, these results indicate that an excessively oxidizing environment, such as that encounter in *E. coli* SHuffle T7 strain, leads to the formation of CdeC aggregates probably as a result of unspecific cysteine bridges due to the high number of consecutive cysteines in the sequence of CdeC, suggesting that the self-assembly properties of CdeC depend on a subtle redox equilibrium.

**Fig. 4.**
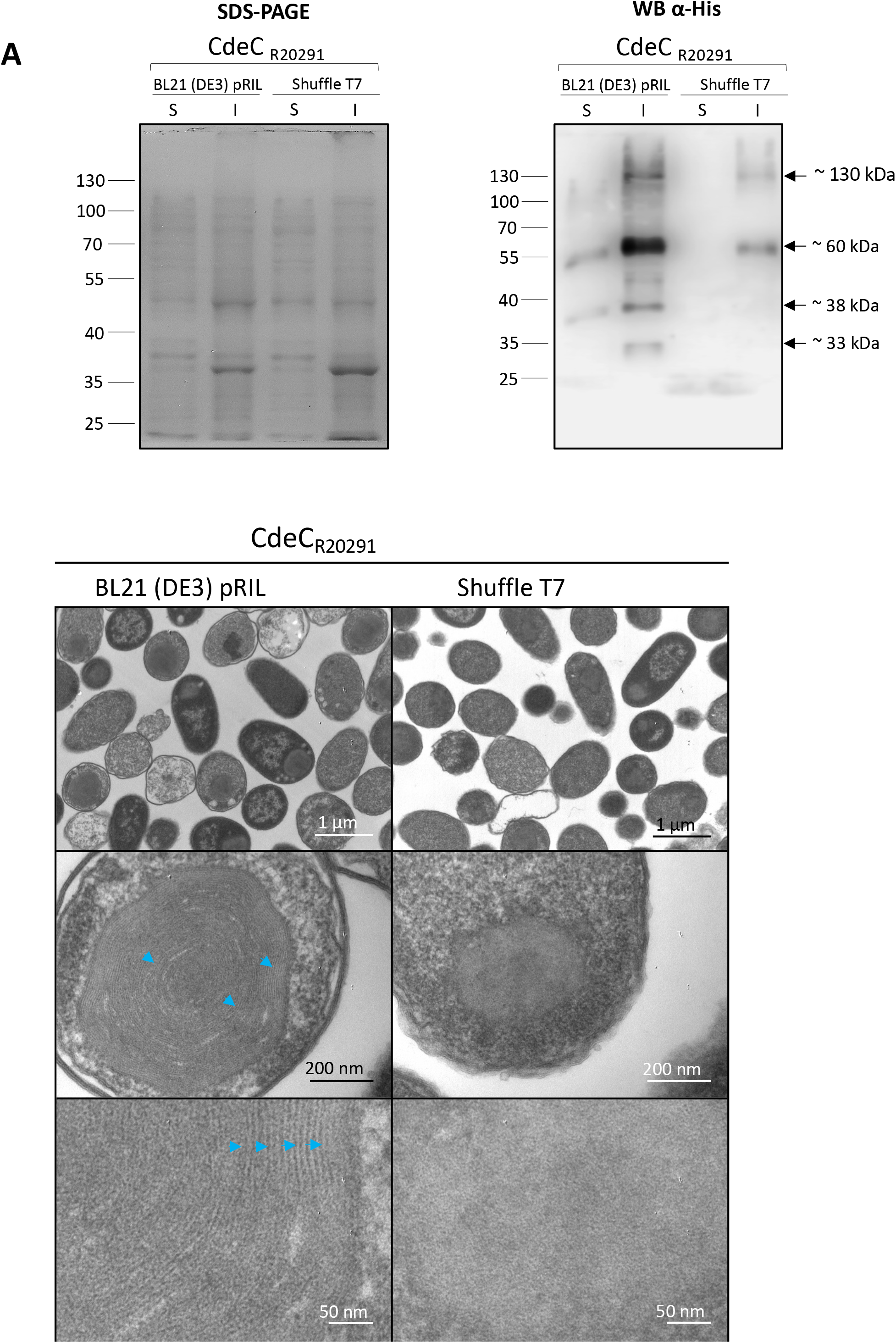
Effect of recombinant strain type in the soluble expression and structural organization of CdeC. A) Recombinant CdeC expression in *E. coli* BL21 (DE3) pRIL and SHuffle T7 strains carrying pARR19 were induced with 0.5 mM IPTG for 16 h at 37°C. Cells were collected and lysed in soluble (S) and insoluble (I) lysis buffer, electrophoresed, and analyzed by Western blot as described in the Methods section. Each lane was loaded with 2 μg of protein lysate. Molecular mass (kDa) markers are indicated at the *left side* of the panels, and molecular mass of the detected His-tagged immunoreactive bands are indicated at the *right side* of the panels. B) Thin sections of *E. coli BL21* (DE3) pRIL and *E. coli* SHuffle T7 expressing CdeC from *C. difficile* R20291 were analyzed by transmission electron microscopy as described in the Method section. Representative micrographs of several *E. coli* cells are shown in the upper panel. The middle panel shows selected individual cells, and the lower panel displays a magnified view of the thin section of inclusion bodies inside *E. coli* BL21 (DE3) pRIL or SHuffle T7. When corresponding, lamellae-like formation in CdeC is indicated with blue arrowheads.

### Effect of DTT on the ultrastructure of CdeC inclusion bodies

Due to the formation of stable and high density, CdeC inclusion bodies can be isolated from cell lysates by differential centrifugation, providing fast, robust, and hence cost-efficient protocols to obtain large amounts of relatively pure protein. We first aimed to test whether CdeC inclusion bodies could be purified from *E. coli* lysates. For this, *E. coli* cells carrying CdeC inclusion bodies were lysed, and the remaining pellet was extracted with Triton X-100, which is a detergent that solubilizes membrane-associated material rather than aggregated proteins. After sequential washes with Triton X-100 and PBS, the pellets resulted in an enrichment of CdeC inclusion bodies that were observed as dark spheres (Figure S4A). Also, all the inclusion bodies presented immunoreactive fluorescence against anti-his antibodies, indicating the accessibility of the 6xHis tag (Fig. S4A). Next, we ask if the purification protocol could affect the ultrastructure of the bodies. For this, a sample of partially purified bodies was analyzed by TEM. TE-micrographs show that the purification protocol did not affect the lamellae-like organization of the inclusion bodies (Figure S4B). Therefore, these results indicate that CdeC inclusion bodies can be partially isolated, retaining their ultrastructural properties and employed for further experiments.

A redox state is the ratio of the interconvertible oxidized and reduced form of a specific redox couple (Schafer & Buettner). The redox state is important since disulfide bond formation involves a reaction between the sulfhydryl (-SH) side chains of two cysteine residues that involves a nucleophilic attack of the S^-^ anion from one sulfhydryl group to a second cysteine, creating the disulfide bond. An oxidative environment will lead to oxidation of sulfhydryl groups and the formation of a cysteine bridge, while a reducing environment will generate the disruption of cysteine bridges (Hatahet et al 2014). Since we observed that an oxidative environment affected the formation of the lamellae-like structures upon heterologous expression in *E. coli* SHuffle T7, we asked whether increasing the reducing environment would affect the stability of the lamellae-like structures.

Therefore, we selected the reducing agent DTT, which maintains sulfhydryl (-SH) groups in a reduced state, it is effective for reducing the disulfide bridges in proteins and the cross-linker N, N’-bis(acryloyl) cystamine. Phase-contrast micrographs demonstrate that the inclusion bodies retained their phase-bright properties upon treatment with various DTT concentrations (Fig. 5A). Also, the DTT treatment resulted in the transition of the inclusion bodies from spherical forms to brighter amorphous structures (Fig. 5A), suggesting a disruption of the compact structure of the inclusion bodies. To assess if the amorphous structures generated after the DTT treatment resulted in less compact structures and, therefore, more accessible to the anti-His antibody, we perform an immunofluorescence assay of each DTT treatment. The immunofluorescence of the partially purified inclusion bodies revealed that there is a qualitative increase in the fluorescence intensity as the concentration of DTT and the incubation time increases. The highest intensity of fluorescence was observed when treating inclusion bodies with 2 M DTT during two hours of incubation (Fig. 5A). The quantification of the fluorescence intensity of inclusion bodies treated with 0.5, 1, and 2 M DTT for 2 hours showed that the fluorescence intensity of Inclusion bodies increased three times when treated with 2 M DTT compared to inclusion bodies not treated with DTT. In contrast, the fluorescence intensities obtained upon treatment of inclusion bodies with 0.5 and 1 M DTT are lower than those of inclusion bodies not treated with DTT (Fig. 5B).

**Fig. 5.**
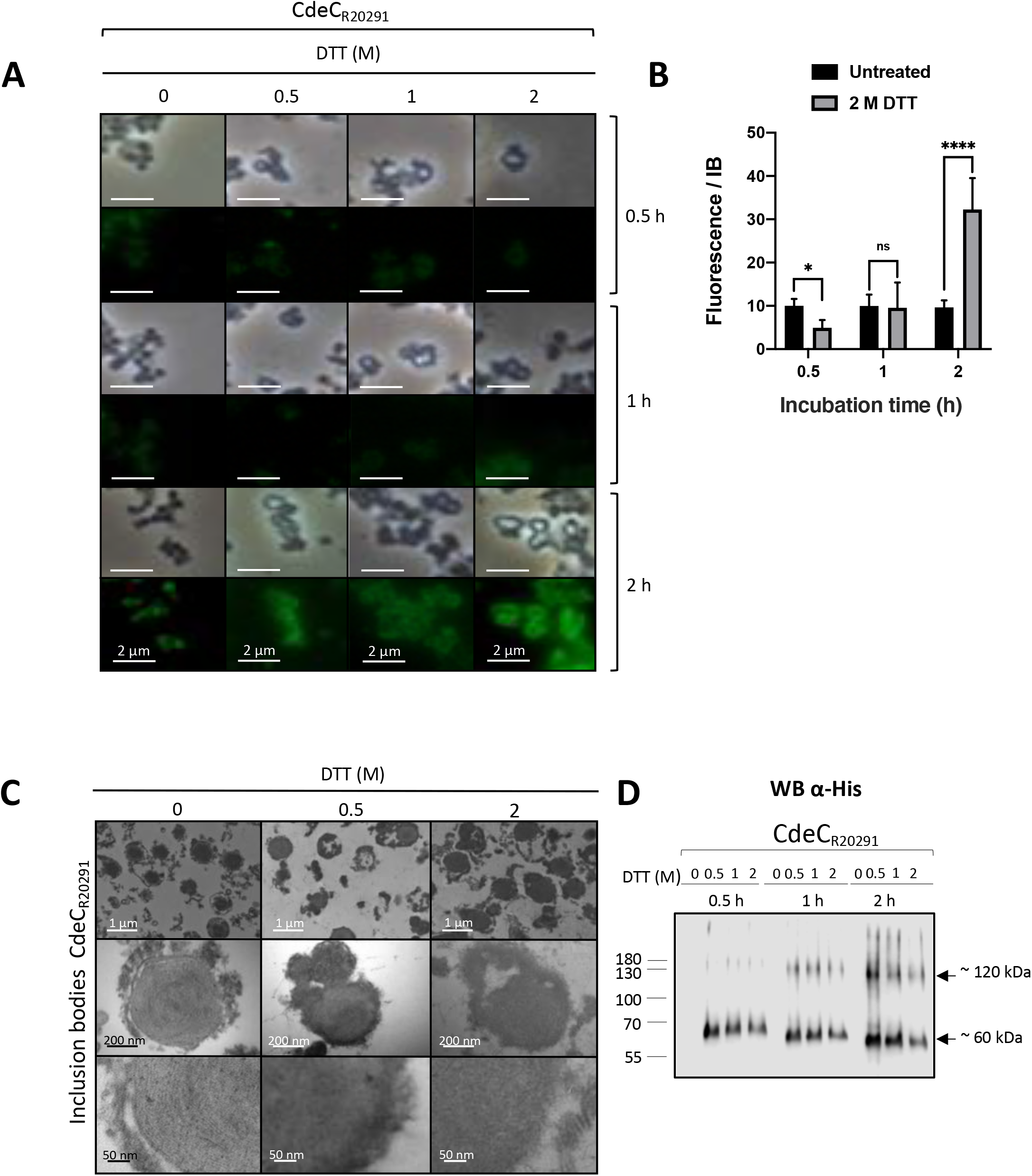
Effect of DTT treatment on the ultrastructure of the inclusion bodies of CdeC. A) The partially purified inclusion bodies were treated with 0.5, 1, or 2 M of DTT for 30, 60, or 120 min at 37°C. After each treatment, the pellet obtained was analyzed by immunofluorescence using as first antibody anti-6His and a secondary antibody conjugated with Alexa-488 fluorophore. B) Quantification of the fluorescence intensity of the inclusion bodies treated with 2 M DTT. The bars in the graph show the mean ± SEM of 10 analyzed inclusion bodies. The statistical significance was determined by a two-way ANOVA analysis using Sidak’s multiple comparisons test. Asterisks denote statistical significance * 0.0236; **** <0.0001; ns (no significance). C) Samples of purified inclusion bodies treated with 0.5 or 2 M DTT during 120 min were analyzed by TEM. The upper panel shows representative micrographs of several inclusion bodies. The lower panel shows a magnified view of the thin section of inclusion bodies. D) Western blot analysis of CdeC inclusion bodies after DTT treatment. The resulting final pellets of each treatment were electrophoresed in 12% SDS-PAGE and His-tagged immunoreactive proteins detected by Western blot analysis as described in the Methods section. Molecular mass (kDa) markers are indicated at the *left side* of the panels, and molecular mass of the detected His-tagged immunoreactive bands are indicated at the *right side* of the panels.

Next, to investigate the effect of DTT treatment in the ultrastructure of the inclusion bodies, samples treated with DTT 0.5 and 2 M during 2 h were observed by TEM. As seen, the characteristic ultrastructure, organized lamellae-like, is mostly lost after the treatment with DTT (Fig. 5C). We also observed that some filamentous fragments of the inclusion bodies are lost after the DTT treatment suggesting those pieces correspond to CdeC aggregates or multimers.

Therefore, to gain more understanding of the effects of DTT on the CdeC inclusion bodies, DTT-treated samples were also resolved by SDS-PAGE prior to immunoblotting with anti-his antibodies. The treatment resulted in the appearance of immunoreactive bands, probably due to the release of monomeric and oligomeric forms of CdeC (Fig. 5D). The inclusion bodies treated with DTT during 30 min resulted in the emergence of a protein band of approximately 60 kDa, suggesting a monomeric form of CdeC. Nevertheless, after 60 min of incubation with DTT, a multimeric form with a molecular weight of approximately 120 kDa is observed in western blot analysis. These higher molecular weight bands may correspond to CdeC dimers or trimers. Interestingly, the longer the incubation time, the immunoreactive bands corresponding to the putative monomeric form run faster and closest to the 55 kDa weight (Fig. 5D). Overall, these results indicate that DTT treatment resulted in the abolition of lamellae-like structures and allowed the release of monomeric and multimeric forms of CdeC from the inclusion bodies to the supernatants.

### CdeC redox-sensitive disordered and residue contact regions

As shown above, CdeC self-assembly properties involved in the formation of the lamellae-like structures depend, at least in part, to the formation of disulfide bridges, and these properties seem to be driven by last 200 amino acids at the C-terminal domain. However, since CdeC contains 37 cysteine residues in its aminoacidic sequence, which might be interacting to drive the formation of the lamellae-like structures, we sought to refine these regions by identifying the redox-sensitive disordered regions (RSDR) using IUPred2A (Meszaros et al 2018). Disordered proteins can mediate protein-protein interactions by recognizing specific partners and undergo a disorder-to-order transition by adopting a more structured conformation (Latysheva et al 2015). The transition can be induced by interactions with other macromolecules or changes in environmental factors, such as pH, temperature, or redox potential (Meszaros et al 2018). The critical sensors built into these redox-regulated proteins are cysteine residues, which can undergo reversible thiol oxidation in response to the oxidation status of the molecular environment (Klomsiri et al 2011). Under reducing conditions, cysteine residues can behave as polar amino acids, most similar to serine, without contributing much to protein stability (Meszaros et al 2018). However, they can also play essential roles in stabilizing the folded conformation by coordinating Zn^2+^ ions under reducing conditions, or by forming disulfide bonds commonly used by extracellular proteins that experience oxidative conditions (Meszaros et al 2018). Using *C. difficile* R20291 CdeC protein, redox-sensitive disordered regions were evaluated (Fig. 6A). We identified the probability of specific regions of being in a disordered or an ordered conformation given its redox state. As seen in Fig. 6A three regions on the protein may have conformations sensitive or dependent to redox state. The first redox-sensitive region is encoded by residues 94 to 109 (CEPCEMDSDECFENKC), which has a predicted CX2CX (6-7)CX4CX(3-4)C sequence-motif. The second redox-sensitive region is encoded by residues 215 to 231 (CETTFEFAVCGERNAEC), exhibits a predicted CX8CX(6-7)C sequence-motif. And the third redox-sensitive region (VDTFSKVCDF), spans from residues 328 to 389, and exhibits a VDXFX3CXF sequence-motif.

**Fig. 6.**
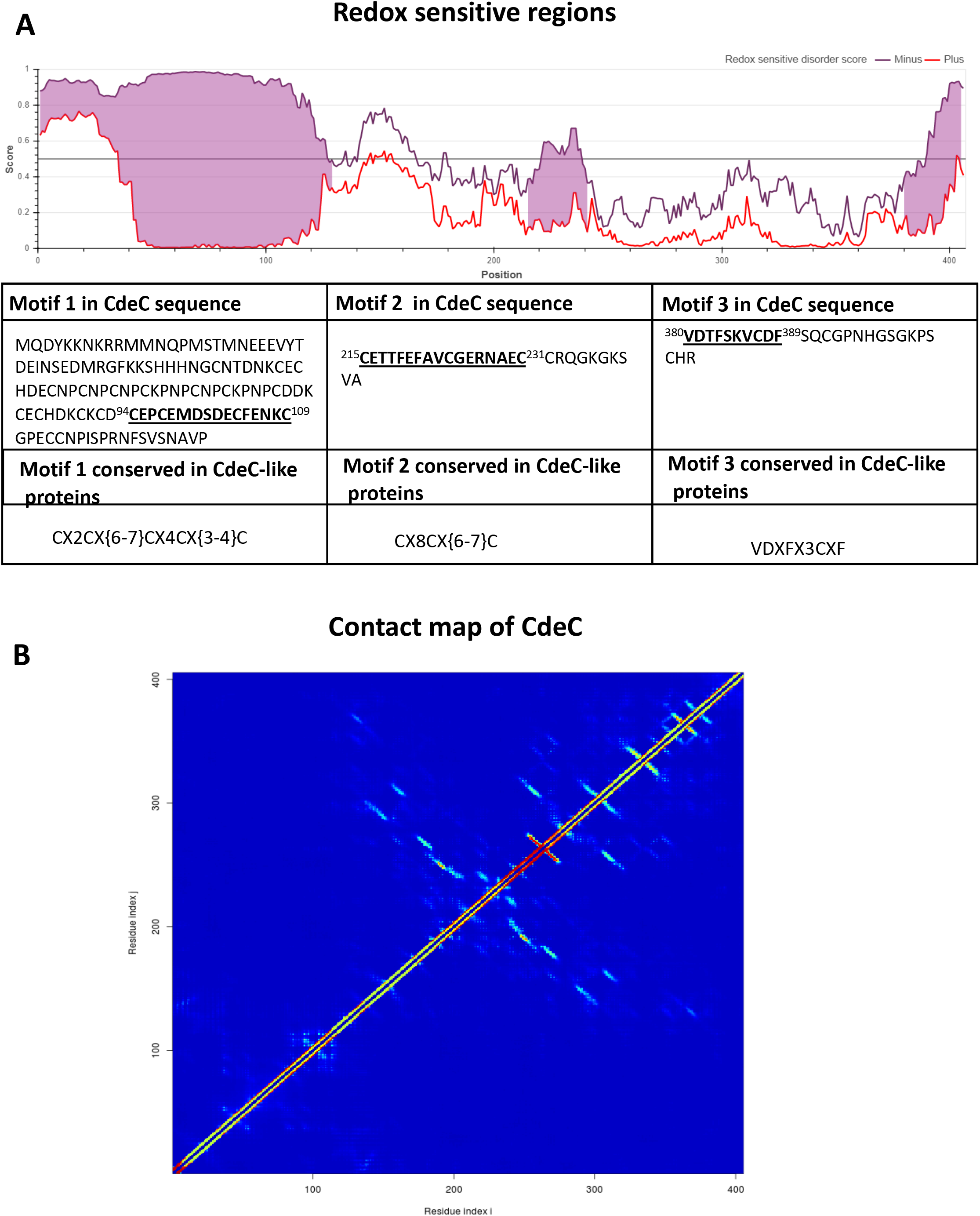
Redox-sensitive disordered regions and residue contact map of CdeC. A) Schematic representation of the prediction of redox-sensitive disordered regions, where the x-axis represents the probability that one position of the protein is disordered (1) or ordered (0). the violet and red lines represent the state that is achieved through cysteine stabilization (redox-plus) and without cysteine stabilization (redox-minus), respectively. When these two profiles are in disagreement, a redox-sensitive region is pinky highlighted. A table attached to the graph with information about the amino acid sequence of the regions and the motifs involved is presented. The motifs are in boldface and underlined in the amino acid sequence. The super index shows the position of the amino acid in the CdeC protein of the *C. difficile* R20291 strain. B) Contact map of CdeC from C. difficile R20291. A heatmap is plotted to show the contact map, interacting interresidues of CdeC, inferred from the amino acid sequence of the protein. The heatmap pattern ranges from red (100% chance of contact) to blue (0% chance of contact). The prediction was made in the web server DeepMetaPSICOV 1.0(Kandathil et al 2019)

To further analyze the putative relevance of the predicted redox-sensitive motifs, and to identify possible pairs of interacting residues, a contact map (a graphical description of interacting partners) was done using the webserver DeepMetaPSICOV 1.0 (Kandathil et al 2019). A color scale based on 100% contact probability (red) to 0% (blue) shows the matrix representation of possible contacts (Fig. 6B). There is a hot-spot region, between position 200 to 250, enriched with putative interacting residues.

### Effect of domain deletions of CdeC on oligomerization and self-assembly properties

The self-assembly of short peptides has been actively studied in recent years (Hu et al 2020). The process is generally driven by specific, spontaneous, and non-covalent chemical interactions. In the case of CdeC, some regions of the protein might have the auto-assembly capacity. Since the redox environment and DTT treatment affects the organized ultrastructure of the inclusion bodies, these organization could depend, at least in part, in the formation of disulfide bridges, suggesting that cysteine residues could be implicated in self-assembly properties. As mentioned before, the sequence of CdeC contains several cysteine residues and sequence repeats (Fig. 7A), but the relevance of these residues is unknown and may be implicated in the organization of lamellae-like structures of CdeC.

**Fig. 7.**
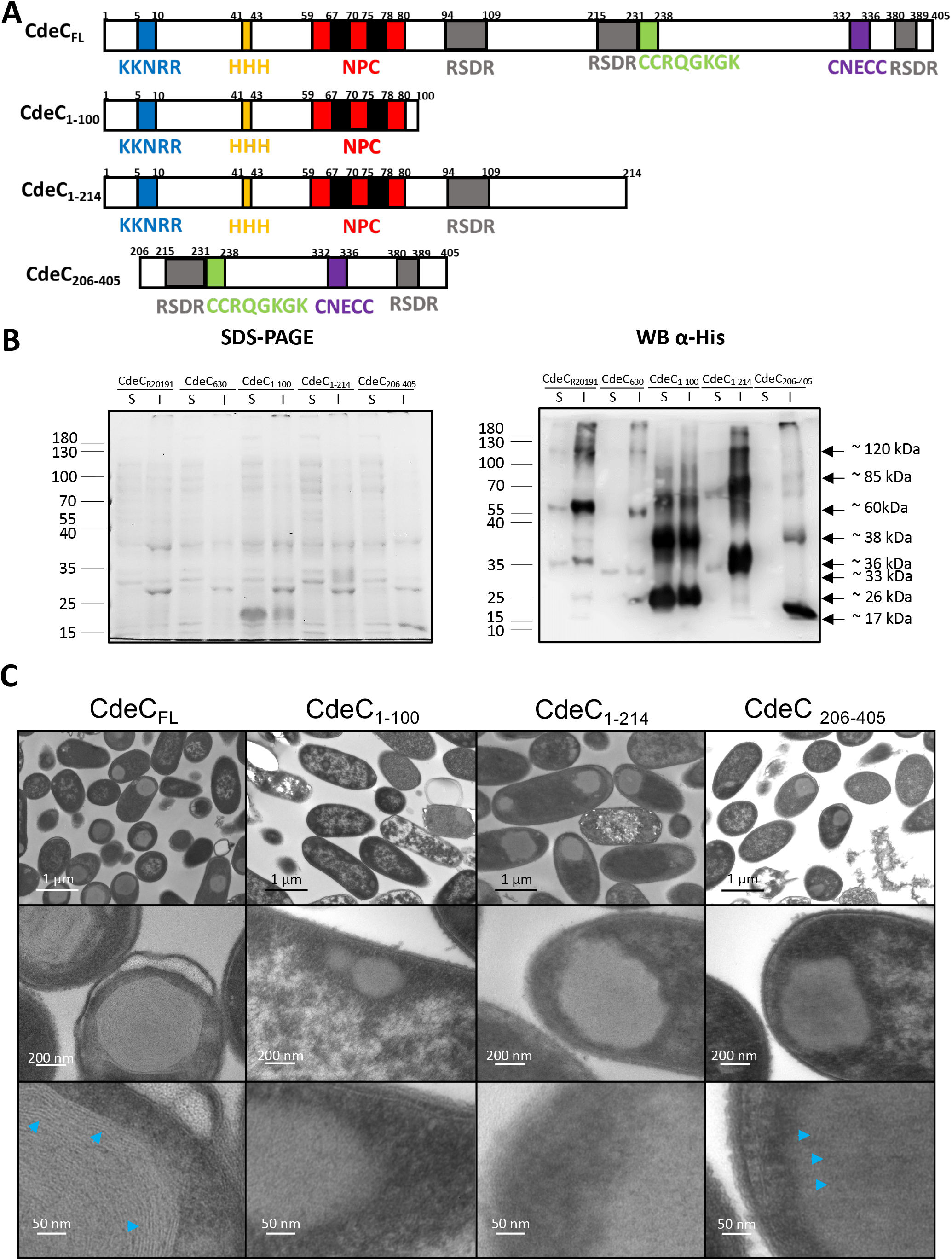
Construction and analysis of CdeC truncations. A) Schematic representation showing the sequence of CdeC (CdeCFL) and the three variants (CdeC1-100, CdeC1-214 and CdeC206-405). B) Recombinant proteins were expressed in *E. coli* BL21 (DE3) pRIL carrying plasmids pARR19, pARR20, pARR7, and pARST1, respectively, and induced with 0.5 mM IPTG for 16 h at 37 °C. The cells were disrupted in soluble and insoluble lysis buffer, electrophoresed in 15% SDS-PAGE, and His-tagged immunoreactive proteins detected by Western blot analysis as described in the Methods section. Each lane was loaded with 2 μg of protein lysate. Molecular mass (kDa) markers are indicated at the *left side* of the panels, and molecular mass of the detected His-tagged immunoreactive bands are indicated at the *right side* of the panels. C) Thin sections of *E. coli* BL21 (DE3) pRIL expressing CdeC from *C. difficile* R20291 and its truncated forms. The sections were analyzed by transmission electron microscopy, as described in the Method section. Representative micrographs of several *E. coli* cells are shown in the upper panel, and the lower panel displays a magnified view of the thin section of inclusion bodies inside *E. coli* BL21 (DE3) pRIL. Blue arrowheads indicate the presence of lamellae-like structures.

Therefore, to define if some of the sequence repeats or some regions in CdeC are prone to aggregation, three different variants were constructed. The first variant denominated CdeC_1-100_ includes the first methionine to the aspartate in position 100. Special features included in this variant are the sequence motifs KKNKRR, HHH, and NPC from the N-terminal domain of the sequence of CdeC (Fig. 7A). The predicted molecular weight of this variant is 11.6 kDa. The second variant, CdeC_1-214_, goes from the initial methionine to the asparagine in position 214. This variant includes the features N-terminal domain KKNKRR, HHH, NPC sequence motifs, and the central region of the sequence of CdeC (Fig. 7A). The redox-sensitive disordered region (RSDR) CEPCEMDSDECFENKC is also present in this variant (Fig. 7A). For this variant, the predicted molecular weight is 22 kDa. The third variant, CdeC_206-405_, starts with the proline in the 206 positions and ends with the arginine 405, the last codon of the CdeC amino acid sequence. This variant includes the CCRQGKGK and CNECC sequence motifs present in the C-terminal domain of CdeC (Fig. 7A)., and two redox-sensitive disordered regions CETTFEFAVCGERNAEC and VDTFSKVCDF (Fig. 7A). The predicted molecular weight of this variant is 23.9 kDa. These three CdeC variants were expressed in *E. coli* BL21 (DE3) pRIL, and *E. coli* lysates obtained analyzed by SDS-PAGE and Western blotting (Fig. 7B). Western blot analysis showed the presence of multimeric forms of CdeC_1-100_ CdeC_1-214_ and CdeC_206-405_ variants. For variant CdeC_1-100_, western blot analysis revealed at least three multimeric forms of apparent molecular weight of 26, 38, and 60 kDa in the soluble and insoluble fractions, suggesting the formation of multimeric forms probably dimers or trimers of CdeC_1-100_. For the CdeC_1-214_ variant, western blot analysis of soluble and insoluble fractions showed the multimeric forms with a molecular weight of approximately 33 (putative monomer) and 70 (putative dimer) kDa in the soluble and insoluble fraction obtained from the *E. coli* lysates. Besides, a multimeric form with an apparent molecular weight of 120 kDa is observed in the insoluble fraction immunoreactive with the anti-His antibody. For variant CdeC_206-405_, western blot analysis revealed at least four multimeric forms with a molecular weight of approximately 17, 38, 60, and 85 kDa in the insoluble fraction. Presumably, the band with an apparent molecular weight of 17 kDa could be the monomeric form of CdeC_206-405_. (Fig. 7A). The higher molecular weight bands may be multimeric forms of CdeC, including dimers, trimers, and tetramers.

Next, to assess the aggregation and self-organization capacity of the different CdeC variants, E. coli cells overexpressing these variants were analyzed by TEM. TEM of the overexpression in *E. coli* BL21 (DE3) pRIL of the three variants (CdeC_1-100_, CdeC_1-214_, and CdeC_206-405_) shows that CdeC_1-214_ and CdeC_206-405_ variants are aggregated in inclusion bodies of similar size (Fig. 7C) These results agree with the Western blot analysis, where most of the protein was found in the insoluble fraction. In contrast, CdeC_1-100_ variant did not seem to form inclusion bodies, and it was found, qualitatively, in the same amounts in soluble and insoluble fractions (Fig. 7B). Furthermore, the inclusion bodies of variants CdeC_1-214_ and CdeC_206-405_ are likely as big as those of CdeCFL but lack the lamella-like structure (Fig. 7C). For the CdeC_206-405_ variant, the inclusion bodies seem to contain discrete laminations (Fig. 7C), but that is not so clear as in CdeCFL. These results may suggest that some of the amino acid residues essential for the CdeC organization may be localized in the C-terminal domain of the protein. Overall, these results demonstrate that the different oligomerization properties of CdeC span the entire amino acid sequence, while the C-terminal domain is involved in self-assembly properties.

## Discussion

The outer-most layer of *C. difficile* spores is critical for understanding the pathogenesis of CDI and for developing novel intervention therapies. Recent work has shown that the composition of this outer layer includes three cysteine-rich proteins (i.e., CdeA, CdeC, and CdeM) (Calderon-Romero et al 2018, Diaz-Gonzalez et al 2015). Genetic studies of CdeC and CdeM have shown although both are essential for the assembly of the exosporium layer, CdeC has pleiotropic roles in the biology of *C. difficile* spores, including spore coat assembly, spore resistance and spore-germination (Barra-Carrasco et al 2013, Calderon-Romero et al 2018). However, the underlying mechanism that drives CdeC- and/or CdeM-mediated assembly of the outer-most exosporium layer remains unclear. In this work, we begin dissecting the underlying molecular basis that drives exosporium-assembly of *C. difficile* spores by demonstrating that although CdeA, CdeC, and CdeM oligomerize upon heterologous expression in *E. coli*, only CdeC exhibits self-assembly properties that lead to the formation of inclusion bodies with lamellae-like structures.

The first major finding of this work was that the cysteine-rich proteins present in the exosporium of *C. difficile*, CdeA, CdeC, and CdeM can form multimers. Upon heterologous expression in *E. coli,* all three proteins (i.e., CdeA, CdeC, and CdeM) are detected not only as monomers but also as higher immunoreactive species that are stable to denaturation conditions (Fig. 1A), mostly in the insoluble fraction. The immunoreactive bands presented a shift in migration, always running slower than expected. This shift in migration of the cysteine-rich proteins could be attributed to the absence of DTT and 2-mercaptoethanol during protein migration, which could contribute to re-oxidation cysteine-residues. These multimeric forms have also been observed in cysteine-rich proteins of the exosporium layer of spores of the *Bacillus anthracis/cereus/thuringensis* group, such as ExsY (Terry et al 2017); similarly, cysteine-rich proteins of the crust layer of *B. subtilis* spores, such as CotY, also have been observed to form stable oligomers (Jiang et al 2015). Two oligomeric forms of 19.4 and ~120 kDa were observed for ExsY protein (Terry et al 2017). While the CotY protein (~19 kDa), a high molecular weight oligomer >80 kDa was observed (Jarrad et al 2015). Both ExsY and CotY oligomers, when treated with a reducing agent such as DTT and heat, are disassembled into their monomeric form (Jiang et al 2015, Terry et al 2017). The formation of CotY oligomers was previously reported when analyzing spore extracts by Western blot. The presence of multimeric forms of 26, 56, and 78 kDa corresponding to monomer, dimers and trimers, respectively, has been described (Zhang et al 1993). Notably, the recombinant proteins ExsY and CotY produced in *E. coli* self-assembled forming hexameric crystal structures that were characterized by heat stability and endurance against denaturing chemicals such as SDS. The total disruption of those structures was accomplished only when heat and a reducing agent (DTT) were applied (Jiang et al 2015, Terry et al 2017)). In *C. sporogenes,* there was also observed that the cysteine-rich exosporium protein, CsxA, self-assembled into a highly thermally stable structure identical to that of the native exosporium when expressed in *E. coli,* (Janganan et al 2020). Therefore, it is tempting to propose that cysteine rich proteins drive the assembly of the exosporium by building a self-assembled structure that serves as a scaffold for the latter recruitment of other exosporium constituents, and this mechanism may be conserved in Clostridiales.

A second major contribution of this work is the observation of organized inclusion bodies upon CdeC overexpression TEM micrographs showed that inclusion bodies of CdeC exhibited an organized lamellae-like structure (Fig. 2C). Interestingly, we recently demonstrated that overexpression of CdeC in *C. difficile* R20291 resulted in the formation of an aberrant exosporium with a visible disorganized lamellae-like structure (Pizarro-Guajardo et al 2020), suggesting that these lamellae-like structures are due to CdeC. In many cases, assembly is primarily driven by hydrophobic interactions, attenuated by electrostatics with directional specificity imposed by electrostatic, van der Waals, and hydrogen bonding interactions (Perlmutter & Hagan, 2015). Although the precise mechanism that guides the formation of the lamellae-like is unknown, our predicted interaction maps suggest that the C-terminal domain seems to have motifs that contribute to self-organization. This is supported by TEM analysis with the CdeC variants showed that the central and carboxyl regions of the proteins tend to form aggregates but are not able to form the lamellae-like as the de CdeC full length.

Interestingly, the variant CdeC206-405 seemed to form small lamellae-like but not easily distinguishable in our conditions. The use of more sensitive technology, as CryoEM, would help in understanding the structural features of CdeC. One constraint in our study is the utilization of truncated variants of the proteins since this approximation may impair the correct folding of the protein. The high homogeneity of CdeC across *C. difficile* strains, as evidenced by our bioinformatics analysis of *C. difficile* published genomes, where the lower identity at the amino acid level was greater than 90 %. Interestingly, despite the differences at the amino acid level between CdeC of strains R20291 and 630, no differences in the formation of lamellae-like structures and differences in the distance of the interspace of these lamellae-like structures were evidenced, indicating that CdeC’s self-assembly properties are conserved across *C. difficile* strains. It would be interesting to trace the evolution of self-assembled properties of CdeC and answer the question if the most recent ancestor of the CdeC, self-organized in these lamella-like structures.

A major question raised by this work is how the redox environment can affect the formation and stability of the lamellae-like structures in CdeC’s inclusion bodies? Disulfide bridge formation requires an oxidative environment for proper formation; thus, it was somewhat surprising that CdeC expression in Shuffle, with a highly oxidative cytoplasmic environment, lead to the formation of inclusion bodies that lacked the formation of the lamellae-like structures. A plausible explanation could be attributed to the aberrant formation of disulfide bridges within and/or between redox-sensitive regions under a highly oxidative environment. That is, multiple unspecific cysteine bridges formed due to the high number of consecutive cysteines in the CdeC sequence might not be appropriately isomerized by the DsbC chaperon, leading to a protein with an improper folding and therefore unable to form native disulfide bonds with itself of another monomer correctly. It was also surprising to observe that upon treatment of the CdeC inclusion bodies with a reducing agent (i.e., DTT), we observed the disappearance of the lamellae-like structures and the release of monomeric CdeC. However, aggregates visualized by TEM micrographs and high molecular immunoreactive bands were also detectable. These results suggest that strong interactions stabilize the CdeC assembly that goes beyond disulfide bridges, and that might be similar to those observed for the *B. subtilis* crust protein, CotY, where strong denaturing conditions were not able to disrupt CotY crystals (Jiang et al 2015). These results are the first report of the ultrastructure of inclusion bodies formed by cysteine-rich proteins present in the exosporium of *C. difficile;* however, whether this organization is relevant for the organization for exosporium assembly remains unclear and is a matter of current work in our group.

## Acknowledgments

To Paula Salgado for the kindly donation of pETM11. A.R-R. was supported by a Fondo Nacional de Ciencia y Tecnología de Chile Postdoctoral Fellowship (Grant 3180692). This work was supported by grants from Fondo Nacional de Ciencia y Tecnología de Chile (FONDECYT Grant 1151025, 1191601) and by Millennium Science Initiative of the Ministry of Economy, Development, and Tourism to D.P-S.

## Supplementary Figures

**Fig. S1.** Effect of temperature and plasmid of expression on the ultrastructure of CdeC. A) Recombinant proteins were expressed in *E. coli* BL21 (DE3) pRIL carrying the plasmid pARR10. The cells were induced with 0.5 mM IPTG for 16 h at 21°C (left) or 37°C (right) in LBG. After the incubation time, samples were retrieved and prepared for TEM analysis, as described in the methodology. The upper panel shows representative micrographs of several *E. coli* cells. The lower panel shows selected individual cells. B) Recombinant proteins were expressed in *E. coli* BL21 (DE3) pRIL carrying the plasmids pARR19 and pARR10 expressing CdeC cloned in plasmid pET22b (left) or pETM11 (right), respectively. The cells were induced with 0.5 mM IPTG for 16 h at 37°C. Representative micrographs of several *E. coli* cells are shown in the upper panel. The lower panel shows selected individual cells.

**Fig. S2** Sequence alignment of CdeC and its representatives from other species. The amino acids preserved in 100% of the analyzed sequences are highlighted.

**Fig. S3**. Topological comparison of Bayesian (a) phylogenetic reconstructions using BEAST (b) and Maximum Likelihood reconstructions using RAxML. The branch coloration shows the Jaccard index, where 1 (blue) implies 100% similarity in the topology in a clade. All *C. difficile* strains fall into a monophyletic clade where their probability *a posteriori* is 100%. Interestingly, the closest common ancestor of CdeC is found in the ancestor between *C. difficile* and *C. mangenotii,* evidenced by the high sequence identity (61%) and that form in the Bayesian inference a monophyletic clade (probability a posteriori = 94.4%) between all *C. difficile* proteins and the two proteins studied (WP_024622166 and WP_02770250)

**Fig. S4**. Purification of inclusion bodies of CdeC. A) inclusion bodies of CdeC from strain R20291 were purified as described in the Methods section. The partially purified inclusion bodies were adjusted to optical densities of 0.2 at 600 nm for immunofluorescence analysis using as first antibody anti-6His and a secondary antibody conjugated with Alexa-488 fluorophore. In the upper panel, contrast phase of inclusion bodies and the lower panel fluorescence of inclusion bodies. B) After purification, one sample of de CdeC bodies was treated for TEM. The upper panel shows selected individual inclusion bodies, and in the lower panel, a magnification of the thin selection is shown. Blue arrowheads indicate the presence of lammella.

