## Supplementary figures and images for "The cysteine-rich exosporium morphogenetic protein, CdeC, exhibits self-assembly properties that lead to organized inclusion bodies in *Escherichia coli*"

### Fig S1

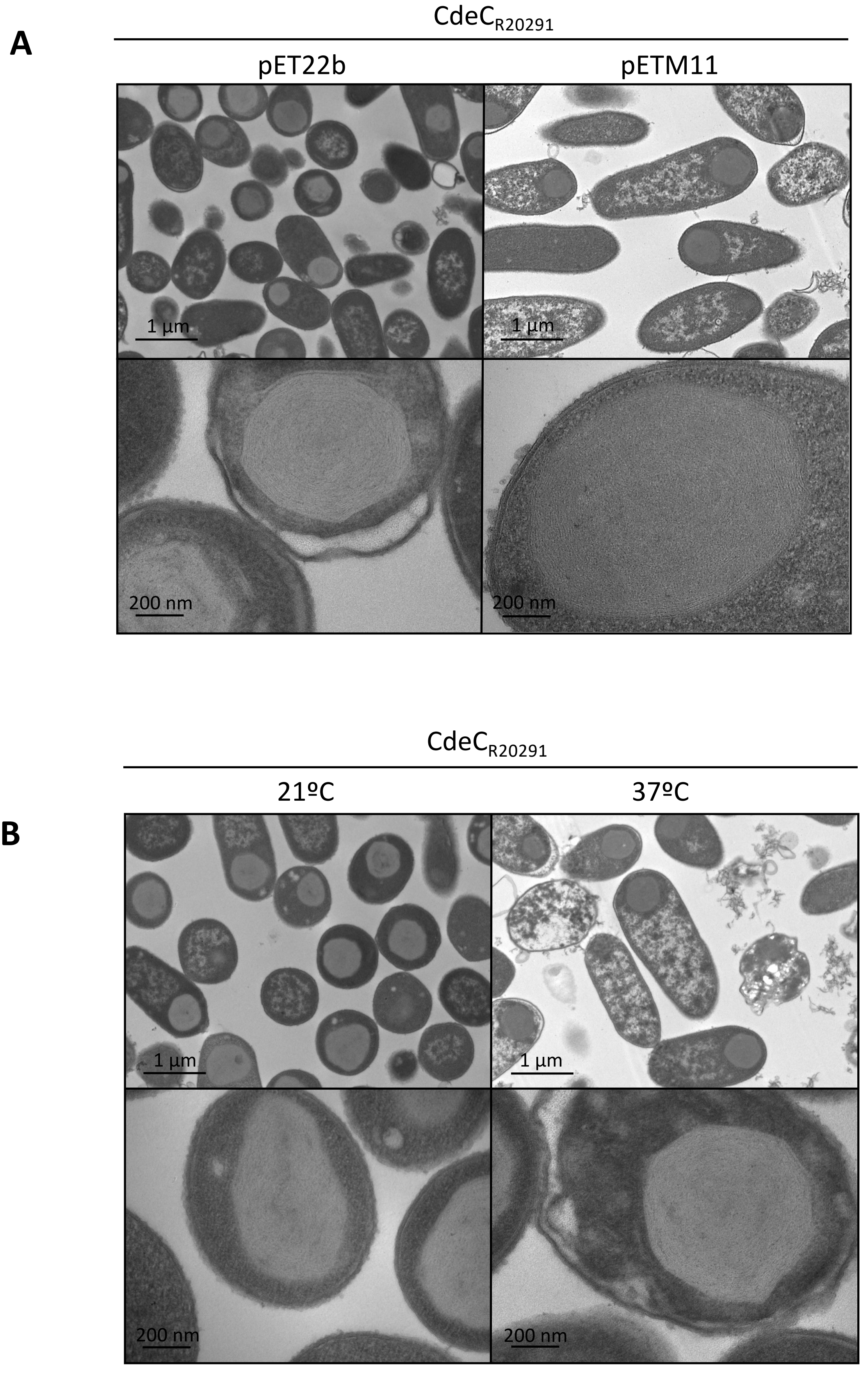

### Fig S2

#### Fig. S2

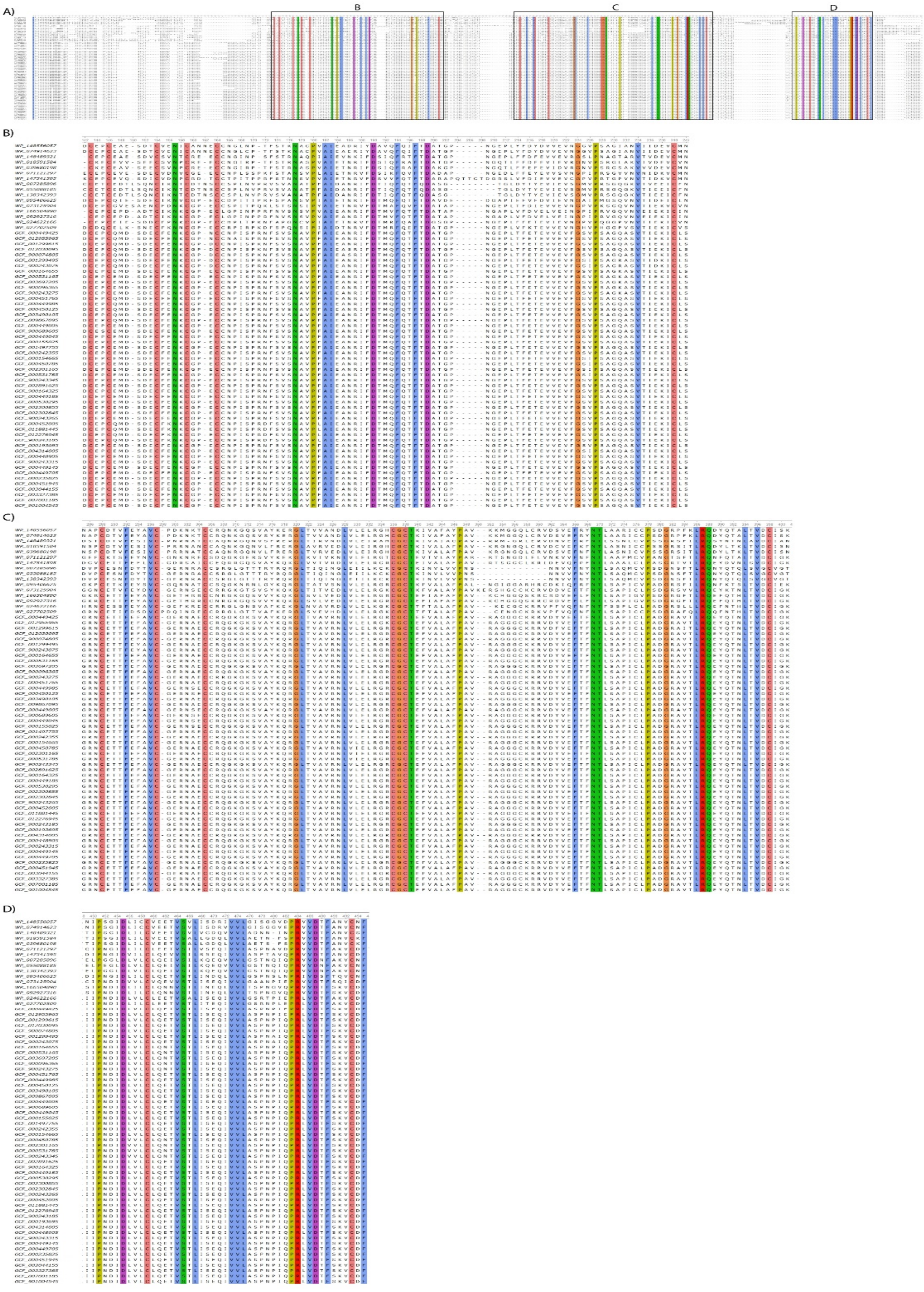

### Fig S3

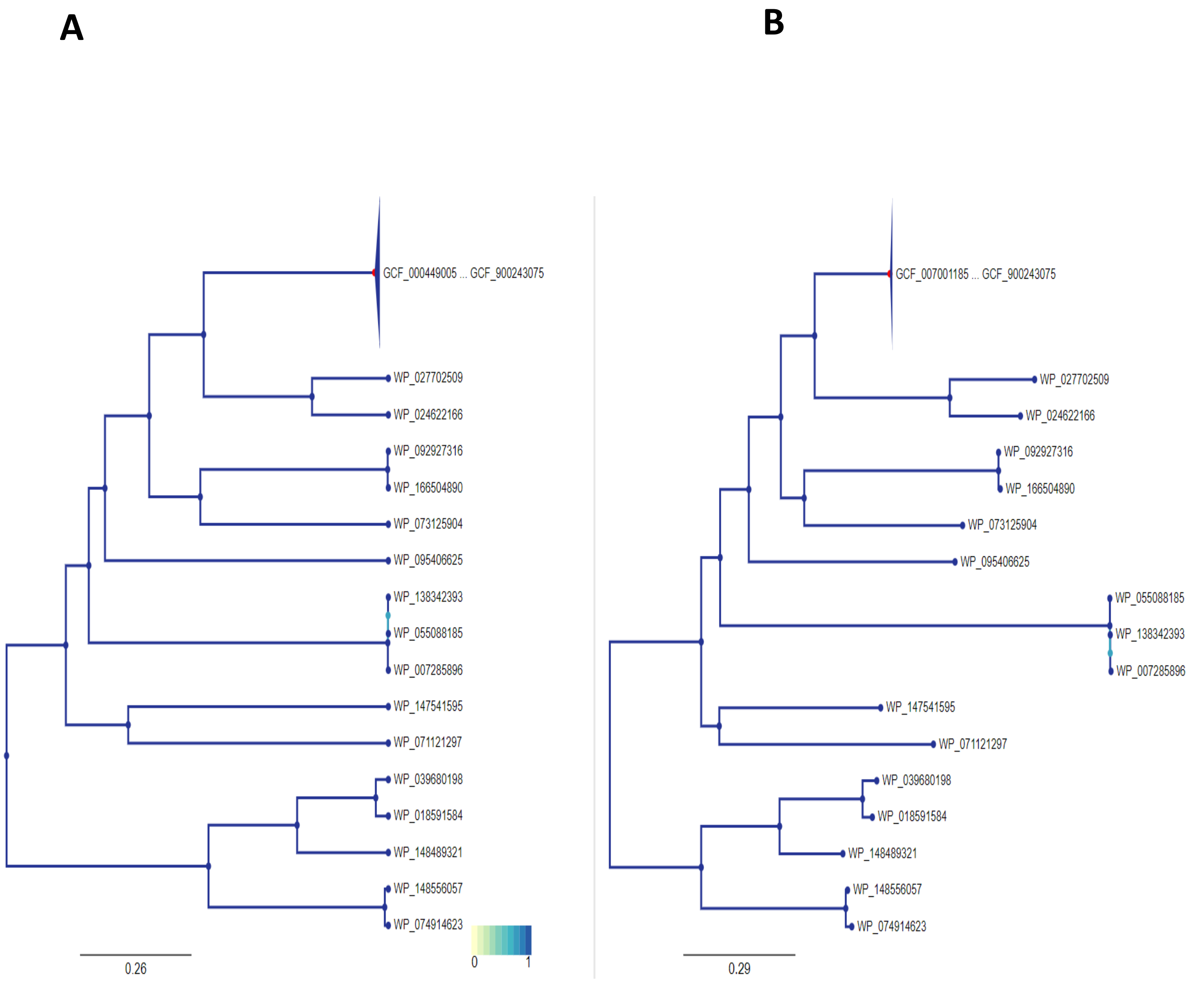

### Fig S4

#### Fig. S4

Purified inclusion bodies of $CdeC_{R20291}$

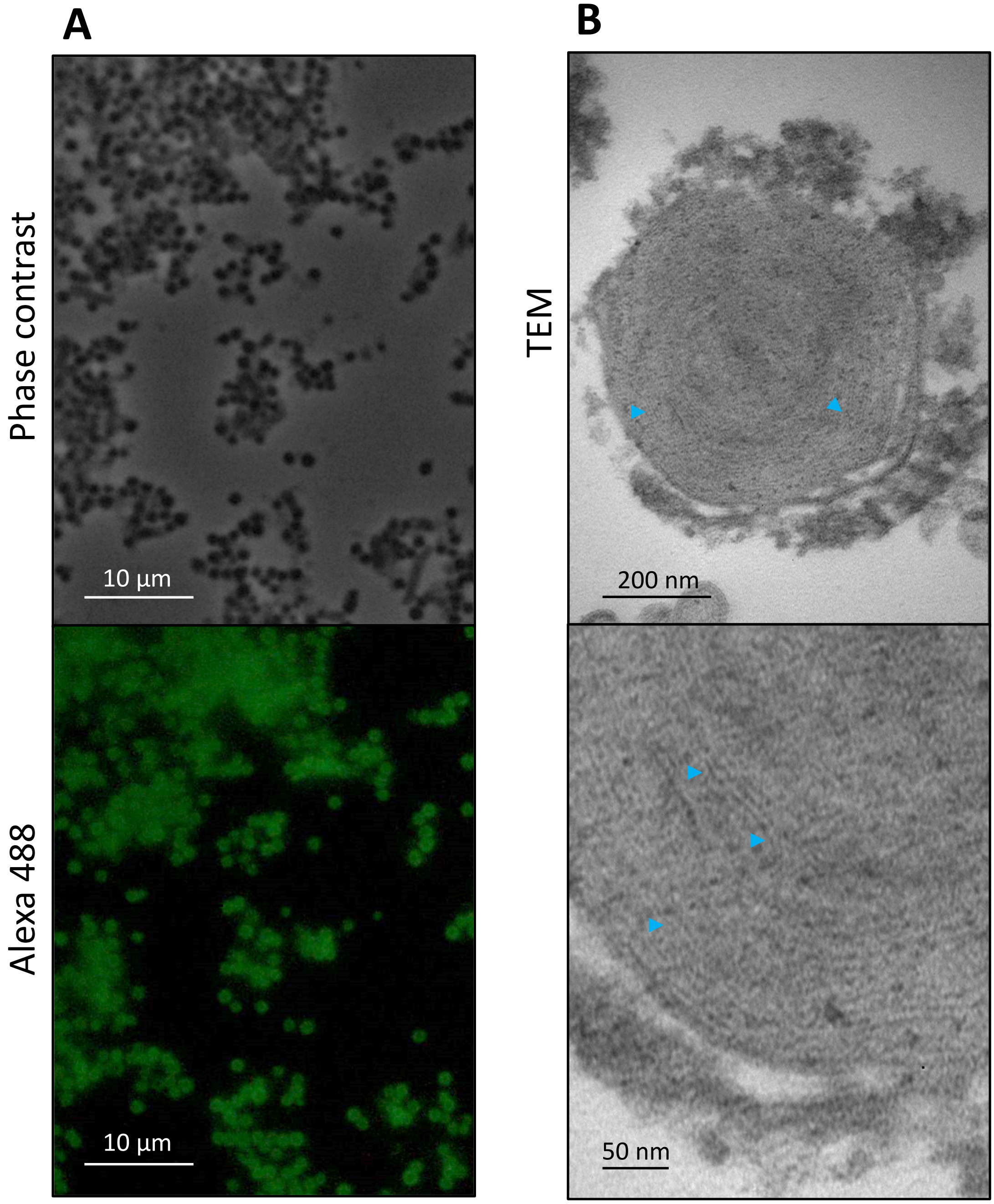
