## Supplementary material for "The cysteine-rich exosporium morphogenetic protein, CdeC, exhibits self-assembly properties that lead to organized inclusion bodies in *Escherichia coli*": Table S1

Table S 1 Plasmids and strain

| **Strains** | Genotype/Relevant characteristic | Reference |
|---|---|---|
| *E. coli* Bl21(DE3) pRIL | $F^-$ *ompT hsdS (rB - mB -) dcm + Tetr gal λ (DE3) endA Hte [argU proL CamR] [argU ileY leuW Strep/SpecR*] / Bl21 strain carrying rare tRNAs codons | Agilent |
| *E. coli* SHuffle T7 | F*' lac, pro, lacIQ / Δ(ara-leu)7697 araD139 fhuA2 lacZ::T7 gene1 Δ(phoA)PvuII phoR ahpC* galE (or U) galK λatt::pNEB3-r1-cDsbC (SpecR, lacIq) ΔtrxB rpsL150(StrR) Δgor Δ(malF*)3 / Derivative of trxB gor suppressor strain SMG96 where its cytoplasmic reductive pathways have been diminished, providing an oxidative environment for disulfide bonded proteins. | NEB |
| *E. coli* DH5α | $F^-$ *endA1 glnV44 thi-1 recA1 relA1 gyrA96 deoR nupG purB20* φ80d*lacZ*ΔM15 Δ(*lacZYA-argF*) U169, hsdR17($r_K^- m_K^+$), $\lambda^-$ | Promega |
| **Plasmids** | | |
| pET22b | The pET-22b (+) vector carries an N-terminal *pelB* signal sequence for potential periplasmic localization, plus optional C-terminal His•Tag® sequence. | Novagen |
| pETM11 | *E. coli* expression vector. Promotor *T7*-Lac, Marker Kanamycin, Tags N-His and C-His, TEV protease cleavage, origin pBR322 | G. Stier EMBL vectors |
| pDP339 | A 1218 pb PCR fragment digested with NdeI and XhoI containing *cdeC* from strain 630, was cloned into N*de*I and X*ho*I sites of pET22b, giving a CdeC-6xHis tag fusion. | (Barra-Carrasco et al 2013) |
| pARR10 | A 1218 pb PCR fragment digested with *Nco*I and *Xho*I containing *cdeC* from strain R20291 ORF, was cloned into *Nco*I and *Xho*I sites of pETM11, giving a CdeC-6xHis tag fusion | This study |
| pARR19 | A 1218 pb PCR fragment digested with N*de*I and X*ho*I containing *cdeC* from strain R20291 ORF, was cloned into N*de*I and X*ho*I sites of pET22b, giving a CdeC-6xHis tag fusion | This study |
| pARR21 | A 492 pb PCR fragment digested with *Nco*I and *Xho*I containing *cdeM* from strain R20291 ORF, was cloned into *Nco*I and *Xho*I sites of pETM11, giving a CdeM-6xHis tag fusion | This study |
| pARR22 | A 305 pb PCR fragment digested with *Nco*I and *Xho*I containing *cdeA* from strain R20291 ORF, was cloned into N*co*I and X*ho*I sites of pETM11, giving a CdeA-6xHis tag fusion | This study |
| pARR20 | A 300 pb PCR fragment digested with N*de*I and X*ho*I containing truncated form M1-D100 of *cdeC* from strain R20291 ORF, was cloned into N*de*I and X*ho*I sites of pET22b, giving a M1-D100-6xHis tag fusion | This study |
| pARR7 | A 642 pb PCR fragment digested with N*de*I and X*ho*I containing truncated form M1-N214 of *cdeC* from strain R20291 ORF, was cloned into N*de*I and X*ho*I sites of pET22b, giving a M1-N214-6xHis tag fusion | This study |
| pARST1 | A 603 pb PCR fragment digested with N*de*I and X*ho*I containing truncated form P206-R405 of *cdeC* from strain R20291 ORF, was cloned into N*de*I and X*ho*I sites of pET22b, giving a P206-R405-6xHis tag fusion | This study |
