## Supplementary material for "The cysteine-rich exosporium morphogenetic protein, CdeC, exhibits self-assembly properties that lead to organized inclusion bodies in *Escherichia coli*": Table S2

Table S 1 List of primers used in this study

| **Primer name** | **Primer sequence [a] 5'-3´** | **Position [b]** | **Gene [c]** | **Use** |
|---|---|---|---|---|
| FP-cdeC-CD0926-N*co*I V2 | CGACCATGGCCATGCAAGATTATAAAAAAAATAAAAGAAGAATG | +3 to +36 | *cdeC*-CD0926 | The forward primer used to clone *cdeC* of in the expression vector pETM11. |
| RP-CdeC-CD0926-X*ho*I V2 | GCGACTCGAGTCTGTGGCAACTTGGCTTTCC | -1195 to-1215 | *cdeC*-CD0926 | The reverse primer used to clone *cdeC* in the expression vector pETM11. |
| FP-CD1067-N*de*I | GACCATATGCAAGATTATAAAAAAAATAAAAGAAGAATGATGAATCAGC | +3 to +45 | *cdeC*-CD0926 | Forward primer used to clone *cdeC* in the expression vector pET22b. |
| RP-CD1067-X*ho*l | GACCTCGAGTCTGTGGCAACTTGGCTTTCCACTTC | -1189 to-1215 | *cdeC*-CD0926 | The reverse primer used to clone *cdeC* in the expression vector pET22b. |
| FP-cdeMpM11 | TCTTTATTTTCAGGGCGCCATGGATATGGAAAATAAAAAATATGCAAATGGTGGTTATTCAGA | +3 to +38 | *cdeM*-CD1478 | The forward primer used to clone *cdeM* in the expression vector pETM11. |
| Rp-cdeMpM11 | TGGTGGTGGTGGTGCTCGAGTTTTCTACAGCAGTTACAATTACATTTATGGCATTTATGG | -453 to -492 | *cdeM*- CD1478 | The forward primer used to clone *cdeM* in the expression vector pETM11. |
| FP-cdeApM11 | TCTTTATTTTCAGGGCGCCATGGTGAAAAATAATAATTTAAATTGTGCTGCTACTAATTGTGCTTATAATACT | +3 to +51 | *cdeA*-CD2262 | The forward primer used to clone *cdeA* in the expression vector pETM11. |
| RP-cdeApM11 | TGGTGGTGGTGGTGCTCGAGTTTCATTTCAAAAGTTTCACAACTTGCATTTCTTTCATTTATATGAAC | -259 to -306 | *cdeA*-CD2262 | The reverse primer used to clone *cdeA* in the expression vector pETM11. |
| FP-M1-D100 CdeC-N*de*I | GACCATATGCAAGATTATAAAAAAAATAAAAGAAGAATGATGAATCAGC | +3 to +45 | *cdeC*-CD0926 | The forward primer used to clone truncated sequences of *cdeC* in the expression vector pET22b, corresponding to coding nucleotides for M1 to D100 of the complete sequence of *cdeC.* |
| RP-M1-D100 CdeC-X*ho*I | GACCTCGAGATCCATTTCACATGGTTCACAATCACATTTAC | -267 to -297 | *cdeC*-CD0926 | The reverse primer used to clone the truncated sequence of *cdeC* in the expression vector pET22b, corresponding to coding nucleotides for M1 to D100 of the complete sequence of *cdeC* |
| FP-M1-N214 CdeC-N*de*I | GACCATATGCAAGATTATAAAAAAAATAAAAGAAGAATGATGAATCAGC | +3 to +45 | *cdeC*-CD0926 | The forward primer used to clone truncated sequences of *cdeC* in the expression vector pET22b, corresponding to coding nucleotides for M1 to N214 of the complete sequence of *cdeC.* |

| | | | | |
|---|---|---|---|---|
| RP-M1-N214 CdeC-*Xho*I | GACCTCGAGGTTTCTTCCTACTATATCTCCTAATGGG | - 610 to -639 | *cdeC*-CD0926 | The reverse primer used to clone truncated sequences of *cdeC* in the expression vector pET22b, corresponding to coding nucleotides for M1 to N214 of the complete sequence of *cdeC*. |
| FP-P206-R405 CdeC-N*de*I | GACCATATGCCATTAGGAGATATAGTAGGAAGAAACTG | +613 to +639 | *cdeC*-CD0926 | The forward primer used to clone truncated sequences of *cdeC* in the expression vector pET22b, corresponding to coding nucleotides for P206 to R405 of the complete sequence of *cdeC*. |
| RP-P206-R405 CdeC-X*ho*I | GACCTCGAGTCTGTGGCAACTTGGCTTTCCACTTC | -1189 to -1215 | *cdeC*-CD0926 | The reverse primer used to clone truncated sequences of *cdeC* in the expression vector pET22b, corresponding to coding nucleotides for P206 to R405 of the complete sequence of *cdeC*. |

[a] Restriction site is marked by an underline.

[B] the nucleotide position number begins from the first codon and refers to the relevant position within the respective gene sequence.

[C] Database: Accession No. FN545816.1
